# Shade Inhibits Cambial Activity in *Populus* Stems by the SPL16/SPL23-Mediated Cytokinin Pathway

**DOI:** 10.1101/2024.09.16.613286

**Authors:** Hongbin Wei, Xingyue Xiao, Jiao Deng, Yi Li, Mengting Luo, Chengshan Zhang, Jinyi Xu, Keming Luo

## Abstract

Trees in natural forests or plantations often encounter neighbor proximity signal that negatively impacts wood production. However, the molecular basis underlying shade-regulation of vascular cambial activity during stem radial growth remains unknown in woody species. Here, we revealed that high stand density and simulated shade (low R/FR ratio) suppress the division and differentiation of cambial cells in poplar stems. A genome-wide screen for *Populus SQUAMOSA PROMOTER BINDING PROTEIN-LIKE* (*SPL*) genes identified that *SPL16* and *SPL23* are preferentially expressed in the phloem and cambium, being downregulated by simulated shade. Knocking out *SPL16/23* impaired cambial activity, whereas phloem-specific overexpression of *SPL16* stimulated cambial proliferation and mitigated the shade-inhibition of cambial activity. Additionally, shade decreased bioactive cytokinin (CK) levels by suppressing the expression of CK biosynthesis genes *IPT5a*, *IPT5b* and *LOG1b* in poplar stems. Molecular and genetic studies reveled that SPL16/23 directly activate *IPT5s/LOG1b* expression to promote CK biosynthesis and cambial activity. Moreover, elevated miR156 expression in shade-treated stems regulated *SPL16/23* at the post-transcriptional level, mediating shade’s effects on cambial activity. Collectively, our findings unravel that the miR156-SPL16/23-IPT5/LOG1-cytokinin pathway operates in the shade-mediated inhibition of cambial activity, providing potential targets for the genetic improvement of shade-tolerant trees.

## Introduction

Wood (also called secondary xylem) is not only an essential reservoir of fixed atmospheric carbon dioxide, but also an abundant source of raw materials for various human applications. The differentiation of vascular cambium into xylem cells in is tightly regulated by internal developmental cues and environmental signals (Begum et al., 2013; Agustí and Blázquez, 2020). In natural forests and plantations, trees frequently encounter high stand density, leading to competition for resources such as water and light, generally resulting in growth decline (Cao et al., 2021). In closed canopies, neighboring plants absorb red (R) light for photosynthesis, whereas reflecting or transmitting far-red (FR) light, resulting in a reduction in the R:FR ratio, a condition described as a shade signal. Extensive studies have shown that, in both herbaceous and woody species, neighbor proximity signals (low R/FR ratios) induce adaptive morphological and physiological responses, such as increased plant height, suppressed branching, reduced leaf size, and upward leaf movement (hyponasty), collectively referred to as shade avoidance syndromes (SAS) (Franklin, 2008; Casal, 2012; Sun et al., 2024). Although the molecular mechanisms underlying the SAS have been well elucidated in the herbaceous model plant *Arabidopsis thaliana*, the effect of shade on cambial activity and xylem development in woody species remains largely unknown.

Plants sense shading and neighbor proximity primarily through phytochromes (phys), the R and FR light photoreceptors, which trigger adaptive responses. The signal transduction pathways mediating shade responses have been studied extensively in annual species such as *A. thaliana* and various cereals (Franklin, 2008; Liu et al., 2021). phyB converts from inactive Pr form to active Pfr upon exposure to R light, while FR light absorption by the active Pfr form reverts it to the inactive Pr state. Studies have shown that active phyB interacts preferentially with phytochrome-interacting factors (PIFs), leading to phosphorylation and subsequent degradation of PIF proteins (e.g. PIF4 and PIF5) or inhibition of transcriptional activity (e.g. PIF7) (Lorrain et al., 2008; Li et al., 2012). Under low R/FR ratios, phyB inactivation releases several PIFs (PIF1/3/4/5/7), which modulate a suite of downstream genes to regulate the SAS. Auxin is a key player in controlling elongation growth, and its biosynthesis and signaling pathway is modulated at several levels by shade singling. PIF proteins directly bind and activate the expression of several *TRYPTOPHAN AMINOTRANSFERASE OF ARABIDOPSIS 1* (*TAA1*) and *YUCCA* genes encoding rate-limiting enzymes in auxin biosynthesis (Sun et al., 2012). The key auxin signaling genes *INDOLE-3-ACETIC ACID INDUCIBLE 19* (*IAA19*) and *IAA29* also act as direct targets of multiple PIFs (Hornitschek et al., 2012), cooperatively promoting elongation growth in response to shade signal. Additionally, increasing evidence suggests that light signaling cross-talks with the microRNA156 (miR156)-mediated aging pathways to mediate various adaptive responses to environmental stresses (Yuan et al., 2023). In *Arabidopsis*, PIFs directly suppress the expression of multiple *MIR156* genes under shade-mimicking conditions, releasing multiple downstream targets *SQUAMOSA-PROMOTER BINDING PROTEIN-LIKE* (*SPL*) family genes to further modulate the SAS (Xie et al., 2017). SPL transcription factors act as pivotal regulators of diverse biological processes in plants, including juvenile-to-adult phase transition, flowering, branching, and stress responses (Wang and Wang, 2015). Notably, it has shown that Arabidopsis plants grown under low light (LL) conditions displayed delayed vegetative phase change, a phenotype distinct from SAS, involving increased miR156/157 expression and repression of *SPL9/13/15* genes in a miR156-independnet manner (Xu et al., 2021). Despite the contrasting traits induced by low R/FR ratios and LL conditions in Arabidopsis, both SAS and LL responses involve the miR156-SPL module, highlighting its importance in regulating plant growth in response to changes in light quality and quantity. To date, studies on functions of the miR156-SPL module have been focused exclusively in vegetative phase change, flowering time, plant architecture, and stress responses. However, its potential role in regulating vascular cambial division and wood formation in woody species remains unexplored.

Wood development initiates with the differentiation of vascular cambium cells into secondary xylem mother cells, followed by cell expansion, secondary wall deposition, and programmed cell death (Sundell et al., 2017). Significant progress has been achieved in understanding key regulators of these developmental stages in tree species (Luo and Li, 2022). Vascular cambium activity is modulated by various phytohormones, including auxin, cytokinin (CK), and gibberellins (GA). Auxin, concentrated in the cambium zone, is crucial for cambial cell proliferation, as demonstrated by reduced wood formation in *Populus* when auxin signaling is disrupted (Nilsson et al., 2008). CK, acting both locally and as a long-distance signal, plays a critical role in cambium regulation. In *Arabidopsis*, simultaneous mutations of four cytokinin biosynthesis genes *ISOPENTYLTRANSFERASE* (*IPT1/3/5/7*) led to a loss of vascular cambium activity and a lack of secondary xylem in hypocotyls (Matsumoto-Kitano et al., 2008). In *Populus* trees, a reduction in CK level by overexpression of a cytokinin catabolic gene *Arabidopsis cytokinin oxidase 2* (*AtCKX2*) specifically in the cambium or phloem led to decreased number of cambium cells (Nieminen et al., 2008; Fu et al., 2021). Conversely, transgenic *Populus* species overexpressing *AtIPT7* or a poplar homolog of *LONELY GUY 1* (*LOG1*), encoding an enzyme in cytokinin biosynthesis, increased endogenous CK concentration and in turn enhanced cambial proliferation and wood formation (Immanen et al., 2016). GA, predominantly accumulated in expanding xylem, also coordinates with auxin and CK to regulate cambial activity, with *Populus* ARF7 integrating GA and auxin signaling to control cambial proliferation (Hu et al., 2022). It is well-established that plants adjust their growth and development in response to light changes through mediating changes in hormone levels and signaling (De Wit et al., 2016; Yang and Li, 2017). However, it remains uncertain whether shade signal could also regulate wood development in trees through hormone-mediated pathways.

Here, to investigate the effects of shade on cambial activity, we cultivated *P. tomentosa* plants under varying stand densities and subjected them to shade-mimicking treatments (reduced R/FR ratios). Both high stand density and simulated shade promoted internode elongation at the expense of radial growth, as indicated by reduced cambial activity and wood formation. We identified miR156-targeted *SPL16* and *SPL23* as positive regulators of cambial activity and demonstrated that shade-mediated inhibition of cambial activity is linked to reduced CK levels. Our molecular and biochemical analyses revealed that SPL16 and SPL23 directly activate CK biosynthetic genes *IPT5a*, *IPT5b*, and *LOG1b*. Furthermore, our findings suggest that shade-induced miR156 contributes to the inhibition of cambial activity. This study provides critical insights into the molecular mechanisms underlying shade-mediated inhibition of cambial activity during secondary growth in poplar.

## Results

### High stand density and simulated shade inhibit cambial activity during secondary growth of *Populus* trees

To examine how high stand density affects stem secondary growth, wild-type (WT) *Populus tomentosa* plants were cultivated in pots and placed in the field, under either low stand density (LSD, 35 cm x 35 cm) or high stand density (HSD, 17 cm x 17 cm) with a closed canopy (Supplemental Figure S1A). Compared to the LSD conditions (R/FR, 1.00), HSD treatment decreased the light intensity by 21.8% and reduced R:FR ratio to 0.62 in between neighboring plants due to canopy shade (Supplemental Figure S1B). After growth for two weeks, the poplars in the HSD group displayed increased plant height, a typical trait of shade avoidance responses (Figure 1A, Supplemental Figure S1C). The 5^th^ to 8^th^ internodes of both LSD and HSD poplars were then collected for anatomical observations stained with toluidine blue. Cross-sections of different internodes revealed that HSD inhibited the stem radial growth of poplar trees compared to those in LSD conditions, as evidenced by a significant reduction in the number of xylem cell layers and cambial cell layers at the 7^th^ internode (Figure 1B-E; Supplemental Figure S1D and E). These results indicate that canopy shade, causing reductions in both light irradiance and R/FR ratios, impairs cambial cell proliferation and secondary growth during wood formation in *Populus*.

**Figure 1.**
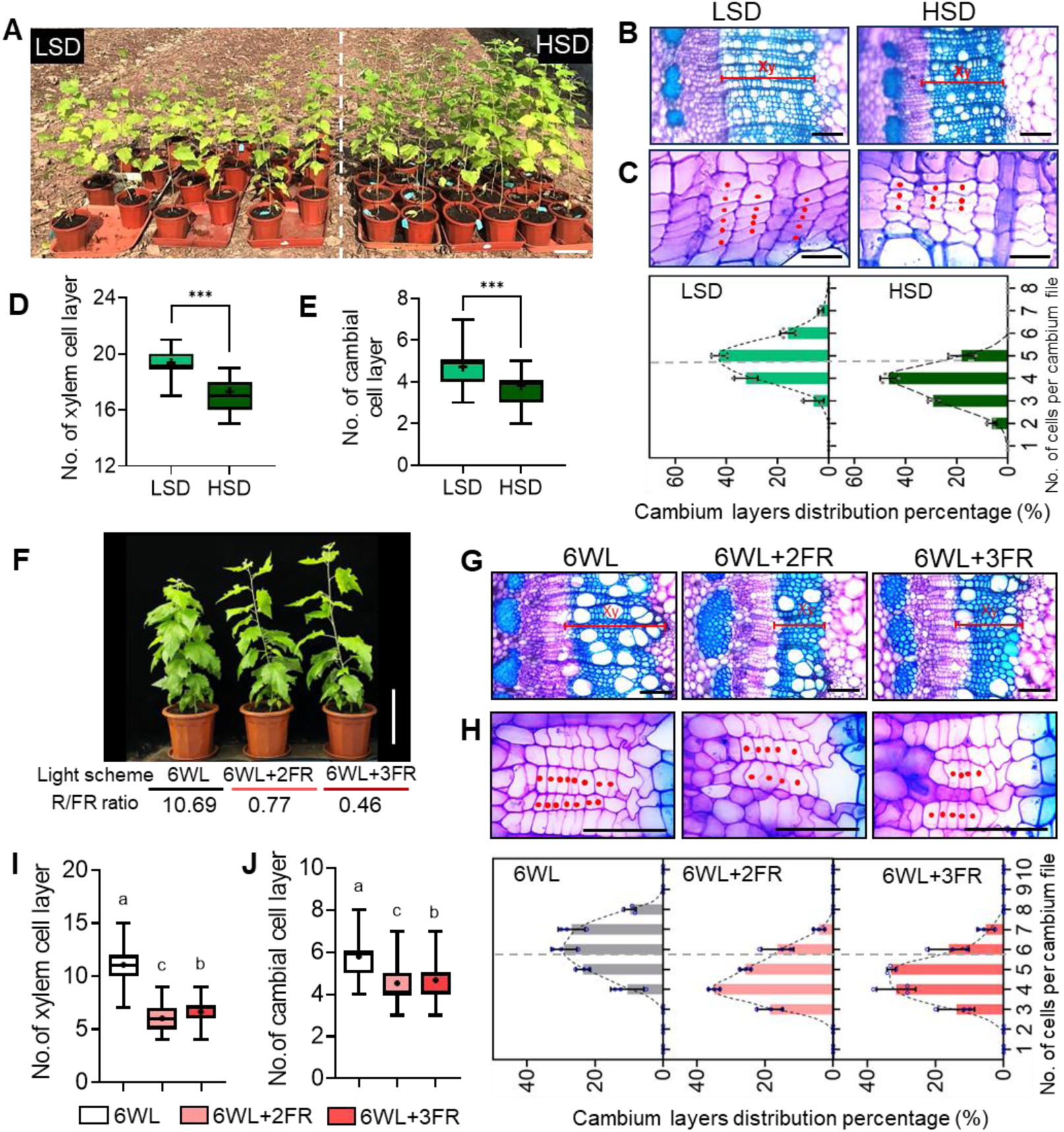
High stand density and simulated shade inhibit cambial activity during stem radial growth in poplar. (**A**) Growth phenotypes of wild type (WT) *P. tomentosa* plants grown under low stand density (LSD, R/FR = 1.00) and high stand density (HSD, R/FR = 0.62) in filed conditions. Bar = 18 cm. (**B**) Xylem phenotypes in the 7^th^ internodes of WT poplars under LSD and HSD. Stem cross-sections were stained with Toluidine blue. Bars = 100 μm. (**C**) Close-ups of cambial cells in cross-sections at the 7^th^ internode (upper panel). Red dots indicate cambial cells in the same file. Bars = 20 μm. Frequency distribution of the number of cells per cambium cell file (lower panel). Values are mean ± SD from three different plants. (**D**) Quantification of the number of cells in each xylem cell file of poplars under LSD and HSD. Values are mean ± SD (n = 100). (**E**) Quantification of the number of cells per cambium cell file of poplars under LSD and HSD. Values are mean ± SD (n = 192). (**F**) Growth phenotypes of poplar WT plants grown under normal light consisting of 6 white lights (6WL, R/FR = 10.69) and simulated shade conditions: 6WL + 2FR (R/FR = 0.77) and 6WL + 3FR (R/FR = 0.46). Bars = 22 cm. (**G**) Toluidine blue-stained anatomical sections of the 7^th^ internodes of WT plants under normal light and simulated shade conditions. Bars = 100 μm. (**H**) Close-ups of cambial cells at the 7^th^ internodes of WT plants under normal light (6WL) and simulated shade (6WL+2FR and 6WL+3FR) conditions (upper panel). Bar = 50 μm. Frequency distribution of the number of cells per cambium cell file (lower panel). Values are mean ± SD from three different plants. (**I**) Quantification of the number of cells in each xylem cell file of shade-treated poplar plants compared with WL controls. Values are mean ± SD (n = 200). (**J**) Quantification of the number of cells per cambium cell file of shade-treated poplar plants compared with WL controls. Values are mean ± SD (n = 360). In the box plots (D, E, I, and J), whiskers indicate the minimum and maximum values, black lines within boxes represent the median values. Different letters indicate significant differences (*P* < 0.05) by one-way ANOVA and LSD test for pairwise comparisons. Asterisks in (D and E) denote significant differences by Student’s *t*-test (***, *P* < 0.001).

To investigate how simulated shade (reduced R/FR ratio only) affects cambial activity and wood formation, *P. tomentosa* plants were subjected to controlled light treatments in the greenhouse. WT poplars were grown under normal light conditions (six white lights, 6WL, R/FR = 10.69) for 10 weeks, then were either maintained under normal light conditions as controls or subjected to WL supplemented with two far-red (FR) lights (6WL+2FR, R/FR = 0.77) or three FR lights (6WL+3FR, R/FR = 0.46) (Supplemental Figure S2A). Reduced R/FR ratios significantly increased plant height, with the greatest increase observed after 8 days of treatment under 6WL+3FR (Figure 1F, Supplemental Figure S2B). Both simulated shade treatments significantly inhibited secondary xylem development, resulting in significant decrease in the number of xylem cell layers compared to the controls (Figure 1G and I). Additionally, fewer cambial cell layers were observed in poplar stems under shade-mimicking conditions (Figure 1H and J). Interestingly, both 6WL+2FR and 6WL+3FR treatments resulted in wider stem diameters associated with the elongation of internodes (Supplemental Figure S2C). Although xylem width was reduced, the increased stem diameter was attributed to enlargement of cortical and pith cells, with a more pronounced effect in the 6WL+3FR treatment than under 6WL+2FR (Supplemental Figure S2D-G). This reflects the dose-dependent effects of R/FR ratios on cell expansion and the resultant internode diameter/length. Given that low R/FR ratio-induced stem elongation involves shade-triggered auxin biosynthesis, we hypothesize that more auxin is accumulated in the cortex and pith to support the elongation growth of internodes.

To better mimic natural canopy shade conditions, *P. tomentosa* plants were subjected to a combined light treatment (4WL+2FR, R/FR = 0.55) that reduced both light intensity by 37.7% and R/FR ratio compared to the normal light condition (6WL) (Supplemental Figure S3A). This 4WL+2FR condition led to a moderate increase in plant height but thinner stems compared to those grown under 6WL, without causing cell expansion in the cortex (Supplemental Figure S3B-D). Cross-sectional analysis of the 7^th^ and 9^th^ internodes confirmed that 4WL+2FR treatment significantly inhibited both xylem development and cambial activity (Supplemental Figure S3E-G). These findings indicate that woody trees sense the reduced R/FR ratio from the neighboring plants to stimulate internode elongation at the expense of cambial division and secondary xylem formation, thereby prioritizing vertical growth to compete more effectively for light. Given that the white light supplemented with 2 FR lights (6WL+2FR, simplified as WL+FR) without reducing photosynthetic irradiance, consistently inhibited cambial activity and wood formation similarly to the high stand density (HSD) or 4WL+2FR conditions, WL+FR was selected for subsequent experiments.

### PtoPIFs may not mediate the inhibition of cambial activity by simulated shade

Extensive studies in *Arabidopsis* have established that phytochrome-interacting factors (PIFs) act as key regulators of the shade avoidance syndromes (SAS) under low R/FR and low-intensity blue light conditions (Casal, 2013). Our recent study revealed that demonstrated that *Populus* PIF3 homologs, PtoPIF3.1/3.2, are involved in modulating shade-induced elongation growth via the conserved PIF3-YUC8-auxin pathway (Sun et al., 2024). Consistent with this, WL+FR (R/FR = 0.77) treatment significantly increased auxin content in the 6^th^ internode, which demonstrated the most evident elongation growth by simulated shade, associated with elevated expression of the auxin biosynthetic gene *YUC8* (Supplemental Figure S4A and B). To determine whether PtoPIF3.1/3.2 are involved in shade-regulation of cambial activity and xylem development, we analyzed stem cross-sections of WT, *Ptopif3.1*, and *Ptopif3.2* knockout mutants. Stem radial growth was inhibited in the mutants compared to WT, as evidenced by a reduced number of cambial cell layers in *Ptopif3.1* and *Ptopif3.2* stems (Supplemental Figure S4C and D). Fewer cells were also observed in the xylem cell files in *Ptopif3.1* and *Ptopif3.2* mutants than in WT (Supplemental Figure S4E-F). These findings suggest that while PtoPIF3.1/3.2 are involved in shade-induced internode elongation, they do not mediate the inhibition of cambial activity and wood formation under shade conditions.

In addition, *Populus* PIF8.1/8.2 paralogs have also been implicated in regulating shade avoidance responses, particularly influencing axillary branch outgrowth acting upstream of the branching regulator *BRANCHED1* (*BRC1*) (Ding et al., 2021; Sun et al., 2024). To assess the involvement of PtoPIF8 in cambial activity, we generated CRISPR/Cas9-mediated *Ptopif8.1* knockout mutant (Supplemental Figure S5A) and compared the secondary growth phenotypes of poplar WT and *Ptopif8.1* mutants. Compared to WT plants, the *Ptopif8.1* mutant showed no significant differences in the number of cambial cell layers or xylem cells layers in stems (Supplemental Figure S5 B-E). These findings suggest that, unlike the PIF-centered regulatory pathways governing the typical SAS traits such as stem elongation and suppressed branching, the shade-mediated inhibition of cambial activity appears to be independent of the photoreceptor-PIF signaling pathways.

### Simulated shade suppresses *SPL16* and *SPL23* expression in the phloem and cambium

In *Arabidopsis*, transcript levels of miR156/157 and their SPL targets are controlled by changes in light intensity and light quality (R/FR ratios). Notably, FR light treatment at the end-of-day (EOD-FR), which simulates canopy shade, represses miR156 expression and alleviates miR156-targeted *SPL* genes to promote multiple shade-avoidance responses such as suppressed branching and accelerated flowering (Xie et al., 2017). Conversely, low light (LL) intensity upregulates miR156 expression and suppresses transcript abundance of *SPLs*, causing a delay in vegetative phase change under limited photosynthetic irradiance (Xu et al., 2021). These observations promoted us to explore whether *Populus* SPL transcription factors respond to simulated shade (low R/FR ratios) and if SPLs may link shading cues to the suppression of cambial activity in poplar stems.

The *Populus* genome contains 26 *SPL* genes, of which 18 *SPLs* are targets of miR156 (Li and Lu, 2014). We first analyzed the expression profiles of poplar *SPL* genes using the AspWood transcriptome database that covers all stages of secondary growth in *Populus tremula* stems (Sundell et al., 2017). Notably, most of the miR156-targeted *SPL*s, including *SPL16/23/25* (Group II), are expressed at higher levels in the phloem and cambium zones (Supplemental Figure S6A and B). Quantitative PCR analysis of those miR156-targeted *SPL* family genes in poplar stem bark tissues under simulated shade conditions (6WL+2R and 6WL+3FR) showed significant downregulation of *SPL16*, *SPL23, SPL20, SPL14, SPL11, SPL13, SPL8*, and *SPL27* (Supplemental Figure S6C). Phylogenetic analysis revealed that *Populus* SPL16 and SPL23 are homologous to *Arabidopsis* orthologs SPL3/4, which serve as important regulators of vegetative phase change and floral meristem identity transition (Wang et al., 2009; Jung et al., 2016; Xu et al., 2016). Interestingly, we re-analyzed the previously reported transcriptome dataset of vascular cambium periodicity in *P. tomentosa* (Chen et al., 2021) and found that *SPL16* and *SPL23* were more highly expressed during the active stage of the cambium than during the dormant stage, suggesting the involvement of *SPL16/23* in regulating cambial activity in poplar. Further expression analysis in different poplar internodes revealed that *SPL16* was preferentially expressed in the 3rd internode, while *SPL23* displayed increasing expression along stem development (Figure 2A). Simulated shade (R/FR = 0.77) treatment consistently suppressed *SPL16* and *SPL23* expression in various developing internodes of poplar trees (Figure 2A).

**Figure 2.**
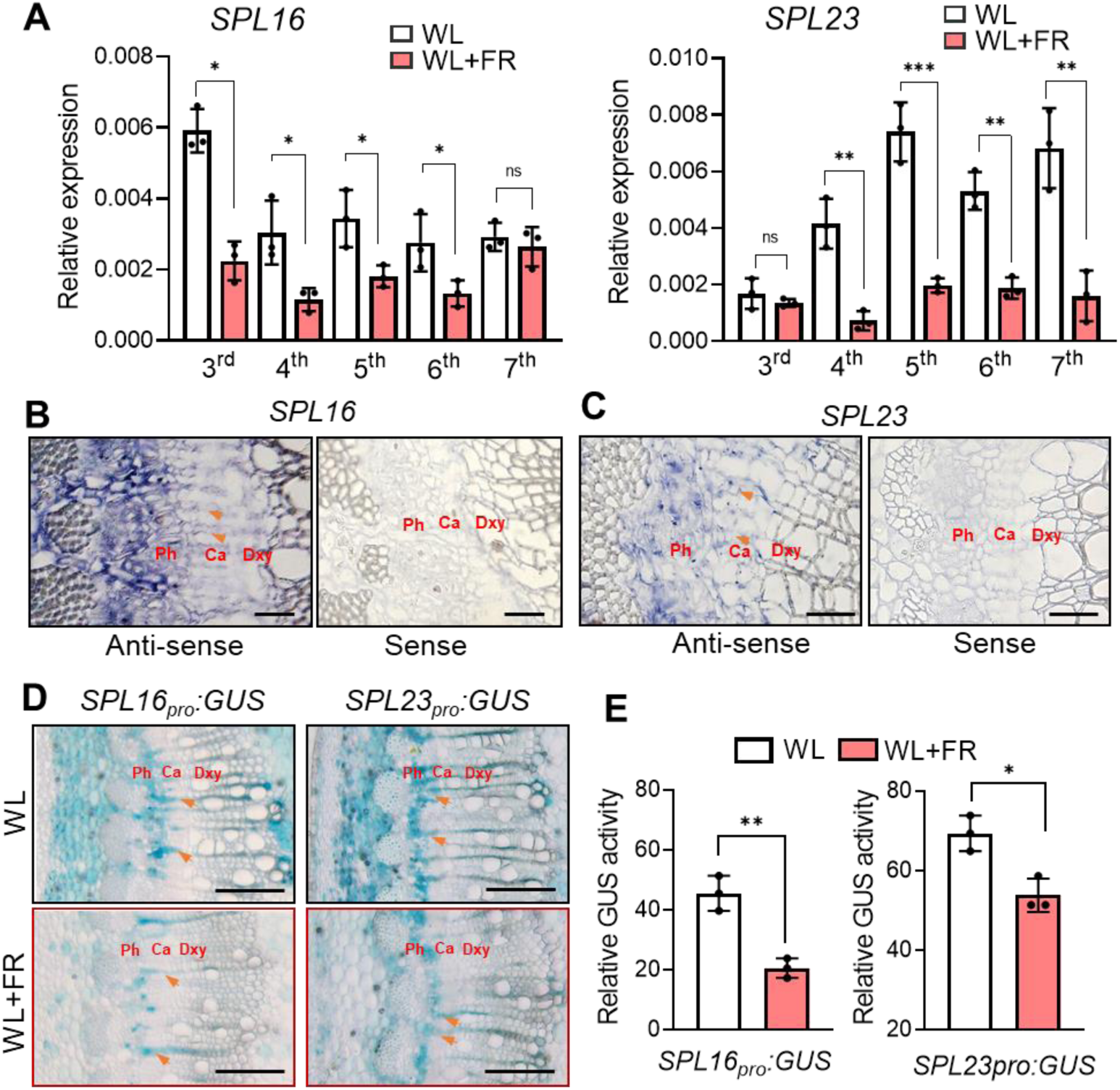
Simulated shade inhibits expression of *SPL16* and *SPL23* in the phloem and cambium zones. (**A**) qPCR analysis of *SPL16* and *SPL23* in developing stems (3^rd^ to 7th counted from the top) of WT poplars under normal light (WL, R/FR = 10.69) and simulated shade (WL+FR, R/FR = 0.77) conditions. *UBQ* was used as the reference gene. Data are mean ± SD from three different plants. (**B** and **C**) *In situ* hybridization analyses of *SPL16* and *SPL23* expression in *P. tomentosa* stems. Cross sections of the 7^th^ internodes from 2-month-old poplar plants were hybridized with antisense and sense probes of *SPL16* and *SPL23*. Blue staining indicates positive *in situ* hybridization signals. Orange arrowheads indicate cambial ray initials. Ph, phloem; Ca, cambium; Dxy, developing xylem. Bars = 50 μm. (**D**) GUS activity in transgenic *Populus* containing *GUS* gene driven by the *SPL16* or *SPL23* promoter. Ten-week-old transgenic plants were grown under WL and WL+FR for 8 days. Orange arrowheads indicate cambial ray initials. Ph, phloem; Ca, cambium; Dxy, developing xylem. Bars = 200 μm. (**E**) Relative GUS activity normalized to protein content in plant extracts. All values shown are mean ± SD from three different plants. Asterisks indicate significant differences by Student’s *t*-test (\**P* < 0.05; \*\**P* < 0.01; ***, *P* < 0.001; ns, not significant).

To verify the expression patterns of *SPL16* and *SPL23* in vascular tissues, we performed RNA *in situ* hybridization. The result confirmed the presence of *SPL16* and *SPL23* transcripts in the secondary phloem and cambial zone, specifically in cambium ray initial cells (Figure 2B and C). Additionally, we generated promoter activity reporter lines harboring the *SPL16_pro_:GUS* or *SPL23_pro_:GUS* construct. GUS staining of cross-sections from these reporter lines confirmed that *SPL16* and *SPL23* are expressed in the phloem and cambial zone of the stem (Figure 2D). After 8 days of simulated shade treatment, we observed reduced GUS signals in these *SPL16_pro_:GUS* and *SPL23_pro_:GUS* transgenic plants. The inhibitory effects of shade on *SPL16* and *SPL23* expression were further confirmed by measuring GUS enzyme activities (Figure 2D and E). These findings indicate that SPL16 and SPL23 might act as positive regulators of cambium activity during secondary growth in poplar.

### Knockout of *SPL16/23* impairs cambial activity while elevated *SPL16* expression mitigates the shade-inhibition of cambial activity

To investigate the roles of *SPL16* and *SPL*23 in regulating cambial activity, we analyzed cambial phenotypes and shade responsiveness in *P. tomentosa spl16* and *spl23* single mutants (Supplemental Figure S7A). WT and transgenic poplars were grown under normal light (WL, R/FR = 10.67) for 6 weeks and subjected to simulated shade treatment (WL+FR, R/FR = 0.77) for 14 days. The *spl16* and *spl23* mutants were slightly taller than WT under WL, but displayed greater increases in plant height compared to WT plants after WL+FR treatment (Supplemental Figure S7B and C). We then examined the cambium phenotypes at the 7^th^ internode of WT and mutant plants under both WL and WL+FR conditions. Cytological observation showed that the number of cambial cell layers was only moderately reduced in *spl16* or *spl23* stems compared to WT under normal WL conditions (Supplemental Figure S7D and E). However, the *spl16* and *spl23* mutants exhibited a more significant reduction in cambial cells compared to WT plants under shade-mimicking conditions (Supplemental Figure S7D and E). Consistently, fewer xylem cell layers were observed in the *spl16* and *spl23* mutants compared to WT, with the most significant reductions occurring in these mutants under shade conditions (Supplemental Figure S7F).

Due to the potential functional redundancy of *SPL16* and *SPL23*, we further examined the cambial activity of *spl16/23* double mutants (L1 and L2) (Supplemental Figure S8A). In accordance with the shade responses of single mutants, the *spl16/23* double mutants demonstrated greater increase in plant height compared to WT plants under shade treatment for 14 days (Supplemental Figure S8B and C). Under normal WL conditions, these *spl16/23* mutants displayed impaired anticlinal division of cambial cells compared to WT plants, as evidenced by fewer cambial cells layers. Simulated shade (low R/FR) exerted even more adverse effects on cambial activity in *spl16/23* mutants than in WT plants (Figure 3A and B). Xylem development was similarly suppressed, with a more marked decrease in xylem cell layers in the *spl16/23* mutants than in WT plants (Figure 3C and D). These results suggest that SPL16 and SPL23 act cooperatively to regulate cambial proliferation and wood formation in *P. tomentosa*. Importantly, our results demonstrated that the cambium division phenotypes resulting from knockout of *SPL16/23* closely resembled those of WT plants subjected to simulated shade, highlighting the critical role of these SPL transcription factors in shade-induced cambial suppression.

**Figure 3.**
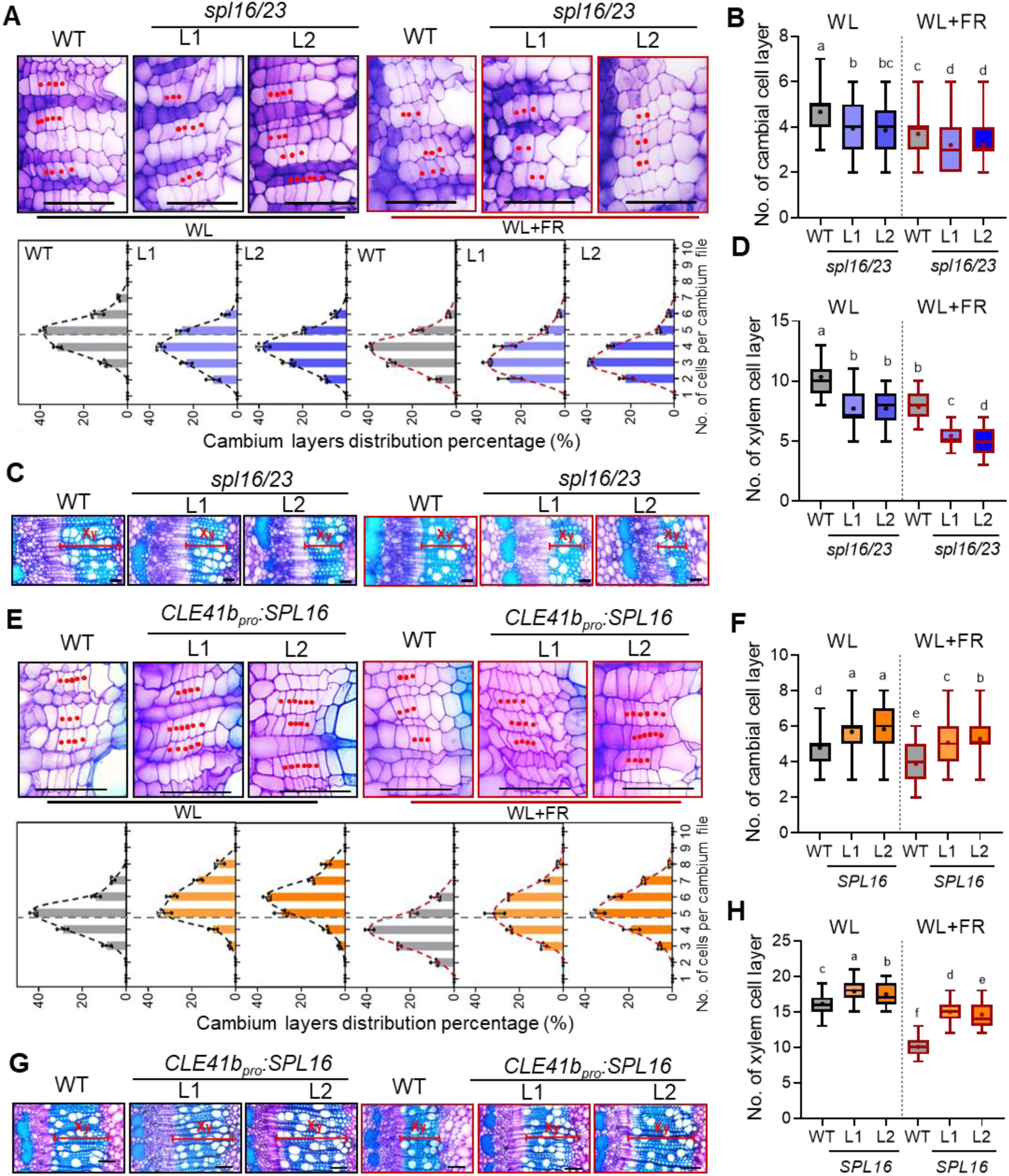
**SPL16 and SPL23 mediate shade-inhibition of vascular cambial activity.** (**A**) Cambial phenotypes of *P. tomentosa* WT and *spl16/23* (L1, L2) mutants grown under normal light (WL, R/FR = 10.67) and simulated shade (WL+FR, R/FR = 0.77) conditions (upper panel). Cross-sections of the 7^th^ internodes from 8-week-old WT and mutant plants were stained with toluidine blue. Red dots denote cambium cells. Bars = 50 μm. Frequency distribution of cell numbers per cambium cell files in the 7^th^ internode of WT and *spl16/23* mutant lines (lower panel). Values are mean ± SD from three different plants. (**B**) Quantification of cambium cell layers in the 7^th^ internode of WT and *spl16/23* mutants. Values are mean ± SD (n = 224). (**C**) The xylem phenotypes at the 7^th^ internodes of WT and *spl16/23* (L1, L2) mutants under WL and WL+FR conditions. Xy, xylem. Bars, 100 μm. (**D**) Quantification of xylem cell layers in the 7^th^ internode of WT and *spl16/23* mutants. Values are mean ± SD (n = 90). (**E**) Cambial phenotypes of WT and *CLE41b_pro_:SPL16* (overexpression of *SPL16* in the phloem) transgenic poplars grown under white light (WL) and simulated shade (WL+FR) conditions (The upper panel). Cross-sections of the 7^th^ internodes from 10-week-old WT and transgenic plants were stained with Toluidine blue. Red dots indicate cambium cells. Bars = 50 μm. Frequency distribution of cell numbers in cambium cell files in the 7^th^ internode of WT and *CLE41b_pro_:SPL16* transgenic plants (lower panel). Values are mean ± SD from three different plants. (**F**) Quantification of cambium cell layers in the 7^th^ internode of WT and *CLE41b_pro_:SPL16* transgenic lines. Values are mean ± SD (n = 240). (**G**) Xylem phenotypes in the 7^th^ internode of WT and *CLE41b_pro_:SPL16* transgenic lines under WL and WL+FR conditions. Xy, xylem. Bars = 100 μm. (**H**) Quantification of xylem cell layers in the 7^th^ internode of WT and *CLE41b_pro_:SPL16* transgenic lines under WL and WL+FR conditions. Values are mean ± SD (n = 200). In all boxplots (B, D, F, and H), whiskers represent minimum and maximum values, black lines indicate median values, and plus symbols denote mean values. Different letters indicate significant differences (*P* < 0.05) by one-way ANOVA and LSD test for pairwise comparisons.

To determine the impact of elevated *SPL16* expression on cambial activity and radial stem growth, we used the phloem-specific *CLE41b* promoter (Fu et al., 2021) to drive *SPL16* expression in the developing secondary phloem of *P. tomentosa* plants. *SPL16* carries complementary sequences to miR156s at the 3’-untranslated region (UTR), thus the *CLE41b_pro_:SPL16* construct is unaffected by miR156s. Among those five independent *CLE41b_pro_:SPL16* transgenic lines generated, L1 and L2 exhibiting high levels of *SPL16* expression were selected for further analyses (Supplemental Figure S9A). Under both WL and WL+FR conditions, *CLE41b_pro_:SPL16* plants showed comparable plant heights to WT plants (Supplemental Figure S9B and C). However, these transgenic plants exhibited enhanced cambial growth and division, confirming SPL16 as a positive regulator of cambial activity (Figure 3E and F). After simulated shade treatment, these *CLE41b_pro_:SPL16* transgenic lines displayed only minor reductions in cambial cell layers compared to WT plants (Figure 3E and F). Additionally, elevated *SPL16* expression in the phloem promoted xylem development and attenuated shade-induced inhibition of xylem differentiation (Figure 3G and H). Taken together, these results indicate that shade inhibits vascular cambial activity during secondary growth through a SPL16/23-mediated regulatory pathway in poplar stems.

### Simulated shade reduces cytokinin (CK) levels by inhibiting CK biosynthetic genes

In *Arabidopsis,* low R/FR ratios inhibit leaf development through auxin-induced cytokinin degradation (Carabelli et al., 2007). Consistent with this, we observed that simulated shade significantly suppressed leaf size in poplar and decreased the contents of iP-type and tZ-type CKs, concomitant with downregulation of CK biosynthesis genes (Supplemental Figure S10A-D). Given the key roles of CK in regulating cambial activity and its specific accumulation in the secondary phloem, we measured CK contents in stem bark tissues of the 7^th^ internode of WT poplars under WL and WL+FR conditions. Results showed that simulated shade also significantly reduced iP and tZ levels in stems (Figure 4A). These results suggest that canopy shade coordinately reduces CK levels in different organs, potentially linking the shade-inhibition of cambial activity to a decline of CK levels in woody tissues.

**Figure 4.**
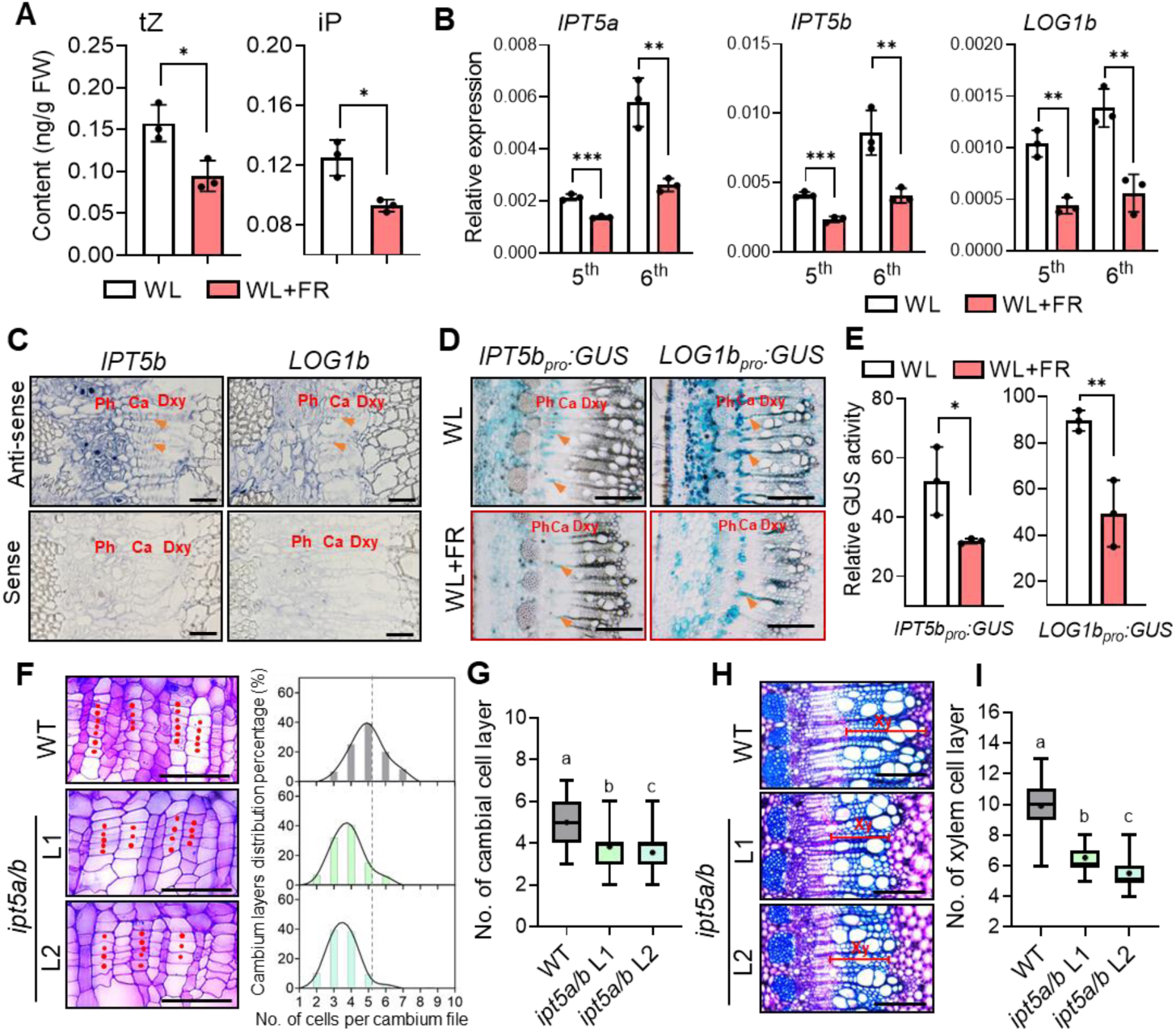
Simulated shade reduces CK levels in stems by inhibiting CK biosynthetic genes. **(A)** Quantification of active CK (iP and tZ) contents in the stem bark of 10-week-old *P. tomentosa* WT plants under normal light (WL, R/FR = 10.67) and simulated shade (WL+FR, R/FR = 0.77) conditions for 8 days. Data are mean ± SD from three different plants. (**B**) qPCR analysis of CK biosynthetic genes *IPT5a*, *IPT5b* and *LOG1b* in stem bark at the 5^th^ and 6^th^ internodes of WT plants under WL and WL+FR for 8 days. Data are mean ± SD from three different plants. (**C**) *In situ* mRNA localization of *IPT5b* and *LOG1b* in the 7^th^ internode of WT plants under WL condition. Stem cross-sections were hybridized with digoxigenin-labeled *IPT5b/LOG1b* antisense RNA probes (upper) or sense probe as controls (lower). Hybridization signals are indicated in blue. Orange arrowheads indicate cambial ray initials. Ph, Phloem; Ca, cambium; Dxy, Developing xylem. Bars = 20 μm. (**D**) GUS staining in stem cross-sections at the 7^th^ internodes of *IPT5b_pro_:GUS* and *LOG1b_pro_:GUS* transgenic poplars under WL and WL+FR conditions. Bars = 200 μm. (**E**) Relative GUS activity normalized to protein content in plant extracts. Data are mean ± SD from three different plants. (**F**) Cambial phenotypes of *P. tomentosa* WT and *ipt5a/b* mutants grown under WL conditions (left panel). The 7^th^ internodes of 10-week-old WT and transgenic poplar plants were cross-sectioned, and stained by Toluidine blue. Red dots denote cambium cells. Bars = 50 μm. Frequency distribution of cell numbers in cambium cell files in the 7^th^ internode of WT and *ipt5a/b* mutants (right panel). (**G**) Quantification of cambium cell layers in the 7^th^ internode of WT and *ipt5a/b* mutants. Values are mean ± SD (n = 120). (**H**) Xylem phenotypes of WT and *ipt5a/b* mutants. Xy, xylem. Bars = 200 μm. (**I**) Quantification of xylem cell layers in the 7^th^ internode of WT and *ipt5a/b* mutants (n = 108). Statistical significance: Asterisks denote significant differences by Student’s *t*-test (\**P* < 0.05; \*\**P* < 0.01; ***, *P* < 0.001). Different letters indicate significant differences (*P* < 0.05) by one-way ANOVA and LSD test for pairwise comparisons.

The initial steps in plant CK biosynthesis are mediated by isopentenyltransferase (IPT) enzymes. LONELY GUY (LOG) enzymes convert conjugated CK nucleotides into bioactive nucleobase forms (Kieber and Schaller, 2014). Thus, we surveyed transcript abundances of CK biosynthesis pathway genes (*IPTs* and *LOGs*) using the AspWood dataset. Five of the nine *Populus IPT* genes show expression profiles in stems, with *IPT5a*, *IPT5b* and *IPT3* displaying a transcript peak in the phloem and cambium zones (Supplemental Figure S11A and B). Among the 17 *LOG* family genes in poplar, *LOG1b* and *LOG2a* are specifically expressed in the phloem and cambium, whereas *LOG1a*, *LOG3a/b*, *LOG5/b*, *LOG7a*, and *LOG8a* are more specific to the developing xylem (Supplemental Figure S12A and B). Thus, our study focused on *IPT5a*, *IPT5b*, and *LOG1b* to understand their roles in CK-mediated regulation of cambial activity in poplar stems.

In accordance with the reduced CK levels in stems, simulated shade significantly inhibited expression levels of *IPT5a*, *IPT5b*, and *LOG1b* in stem bark tissues of WT poplar plants, compared to those in WL-grown plants (Figure 4B). RNA *in situ* hybridization confirmed that *IPT5b* and *LOG1b* were preferentially expressed in secondary phloem and cambial ray initials (Figure 4C). To further analyze the expression profiles of *IPT5b* and *LOG1b*, we generated *IPT5b_pro_:GUS* and *LOG1b_pro_:GUS* reporter lines. GUS staining of these transgenic plants verified that *IPT5b* and *LOG1b* are preferentially expressed in the phloem and cambial zones, wherein the *IPT5b/LOG1b* promoter-driven *GUS* expression was significantly inhibited by simulated shade treatment (Figure 4D and E).

Previous studies have shown that a reduction in cytokinin levels via overexpression of a *CKX* gene or disruption of the CK receptors in phloem impaired cambial activity (Nieminen et al., 2008; Fu et al., 2021). These findings align with our findings of inhibited cambial activity and reduced active CK concentrations under simulated shade. To further explore the roles of *Populus IPT5a* and *IPT5b* in cambial activity during wood formation, we generated *ipt5a/b* knockout mutants using four sgRNA-targeted sites in the highly conserved regions of their cDNAs (Supplemental Figure S13A). The *ipt5a/b* mutant lines (L1, L2) exhibited remarkably decreased plant height and leaf size compared to WT plants (Supplemental Figure S13B and C). Cross-sectional analysis of stems revealed that knockout of *IPT5a/b* resulted in decreased cell layers in the cambium zone (Figure 4F and G), and reduced xylem tissue formation (Figure 4H and I). These results collectively confirm that shade negatively regulate cambium activity during secondary growth by decreasing CK concentration in poplar stems (mainly in the secondary phloem).

### Enhanced CK accumulation in the phloem attenuates the shade-inhibition of cambial activity

To investigate the impact of elevated endogenous CK levels in the phloem on shade-regulation of cambial activity, we generated transgenic poplar lines expressing *IPT5b* gene driven by the phloem-specific *CLE41b* promoter. Among seven independent lines, L1 and L2 with higher *IPT5b* expression in the stem bark were selected for further analysis (Supplemental Figure S14A). These *CLE41b_pro_:IPT5b* transgenic lines exhibited normal plant height but larger leaves, suggesting enhanced endogenous CK levels in stems and that CK has also been transported through phloem into leaf blade to stimulate leaf expansion (Supplemental Figure S14B and C), To explore the effects of enhanced *IPT5b* levels on cambial activity, WT and *CLE41b_pro_:IPT5b* transgenic plants were subjected to simulated shade (WL+FR) for two weeks. These transgenic lines showed significantly higher transcript levels of *type-A RR* genes *RR2* and *RR10*, marker genes for CK accumulation in the phloem (Fu et al., 2021), in the stem bark than those in WT plants under WL+FR conditions (Supplemental Figure S14D), indicating that phloem-specific expression of *IPT5b* could maintain high CK levels in the phloem under canopy shade. Correspondingly, both *CLE41b_pro_:IPT5b* transgenic line exhibited increased cambial cell layers compared to WT under WL conditions, and a less reduction in cambial cell layers when subjected to simulated shade (Figure 5A and B). Additionally, xylem development was also significantly enhanced in *CLE41b_pro_:IPT5b* transgenic plants, with only a moderate decrease observed under shade conditions compared to WT (Figure 5C and D).

**Figure 5.**
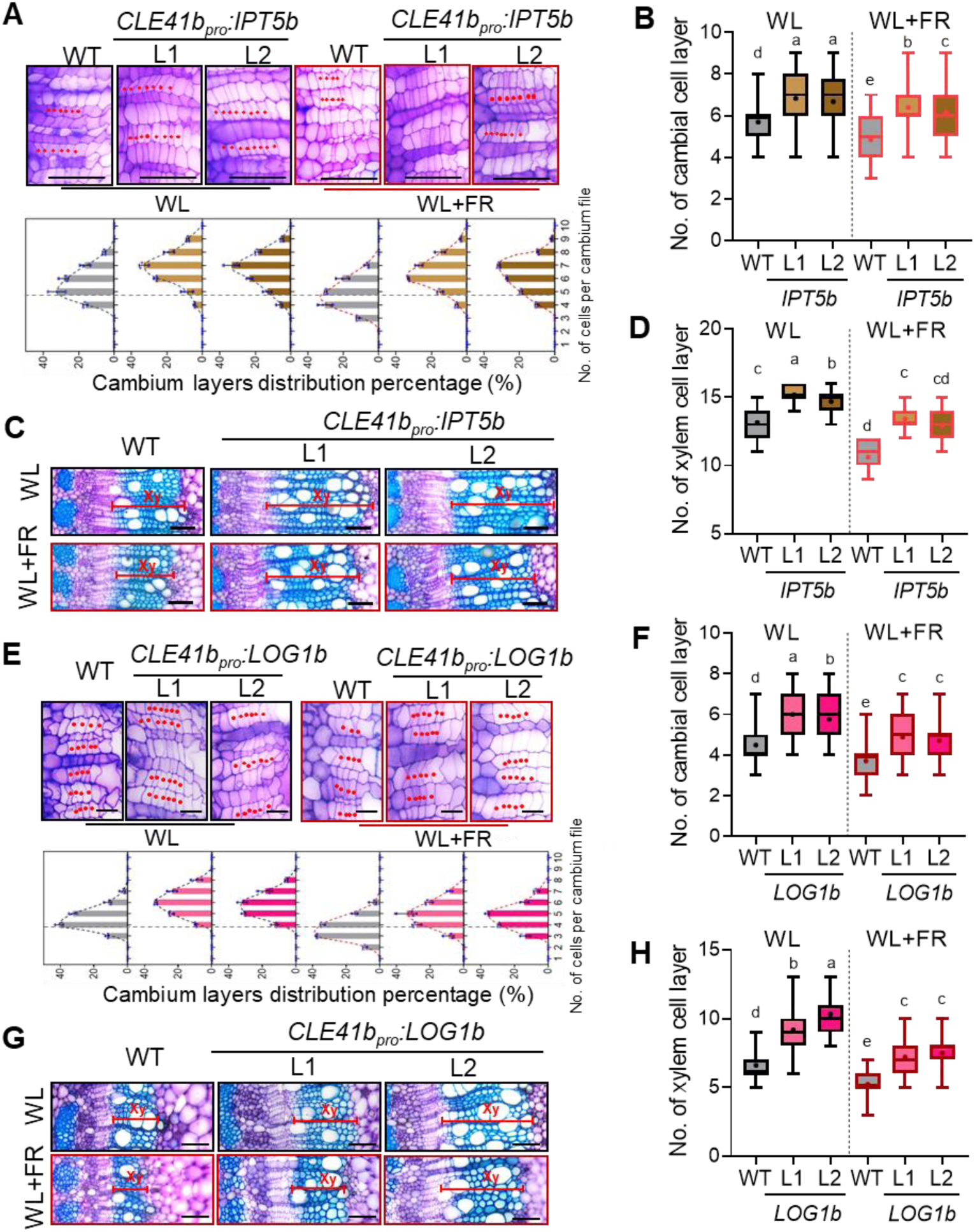
Overexpression of *IPT5b* and *LOG1b* attenuates the shade-mediated inhibition of cambial activity in poplar. (**A**) Cambial phenotypes of *P. tomentosa* WT and *CLE41b_pro_:IPT5b* transgenic plants under white light (WL, R/FR = 10.67) and simulated shade (WL+FR, R/FR = 0.77) conditions (upper panel). The 7^th^ internodes of 10-week-old WT and transgenic poplar plants were cross-sectioned, and stained by Toluidine blue. Red dots denote cambium cells. Bars = 50 μm. Frequency distribution of cell numbers in cambium cell files in the 7^th^ internode of WT and *CLE41b_pro_:IPT5b* transgenic plants (lower panel). Data are mean ± SD from three different plants. (**B**) Quantification of cambium cell layers in the 7^th^ internode of WT and *CLE41b_pro_:IPT5b* lines. Values are mean ± SD (n = 240). (**C**) Xylem phenotypes (stained with Toluidine blue) of WT and *CLE41b_pro_:IPT5b* transgenic plants under WL and WL+FR conditions. Xy, xylem. Bars, 100 μm. (**D**) Quantification of xylem cell layers in the 7^th^ internode of WT and *CLE41b_pro_:IPT5b* transgenic lines. Values are mean ± SD (n = 90). (**E**) Cambial phenotypes of *P. tomentosa* WT and *CLE41b_pro_:LOG1b* transgenic plants under WL and WL+FR conditions (upper panel). The 6^th^ internodes of 10-week-old WT and transgenic poplars were cross-sectioned, and stained by Toluidine blue. Red dots denote cambium cells. Bars = 20 μm. Frequency distribution of cell numbers in cambium cell files in the 6^th^ internode of WT and *CLE41b_pro_:LOG1b* transgenic plants (lower panel). Data are mean ± SD from three different plants. (**F**) Quantification of cambium cell layers in the 6^th^ internode of WT and *CLE41b_pro_:LOG1b* transgenic lines. Values are mean ± SD (n = 240). (**G**) Xylem phenotypes of WT and *CLE41b_pro_:LOG1b* transgenic plants under WL and WL+FR conditions. Xy, xylem. Bars, 100 μm. (**H**) Quantification of xylem cell layers in the 6^th^ internode of WT and *CLE41b_pro_:LOG1b* transgenic lines. Values are mean ± SD (n = 90). Whiskers indicate the minimum and maximum values, black lines within boxes denote the median values, with the means indicated by plus symbols. Different letters indicate significant differences (*P* < 0.05) by one-way ANOVA and LSD test for pairwise comparisons.

To further explore the function of *LOG1b* in cambial development during secondary growth in response to shade, we generated *CLE41b_pro_:LOG1b* transgenic poplar, in which active cytokinin levels (iP and tZ) in the stem were increased by the phloem-specific expression of *LOG1b* (Supplemental Figure S15A and B). Similar to *CLE41b_pro_:IPT5b* transgenic lines, the *CLE41b_pro_:LOG1b* lines (L1, L2) displayed an increased number of cambial cell layers compared to WT plants. Remarkably, the shade-induced inhibition of cambium growth was markedly attenuated in these transgenic lines (Figure 5E and F). Xylem development in the *CLE41b_pro_:LOG1b* plants also mirrored the trends observed in cambial activity under both normal light and simulated shade conditions (Figure 5G and H). These findings validate that *Populus* IPT5b and LOG1b are rate-limiting enzymes of CK biosynthesis pathway involved in prompting cambial activity. Importantly, these results confirm that shade inhibits cambial activity during secondary growth by reducing IPT5/LOG1b-mediated CK levels in the phloem, while enhanced endogenous CK levels antagonize the inhibitory effects of shade on cambial activity in poplar.

### SPL16 and SPL23 regulate CK levels by directly activating *IPT5a*, *IPT5b* and *LOG1b* expression

Given that simulated shade significantly inhibited the expression of *SPL16, SPL23,* and CK biosynthesis pathway genes (*IPT5a*/*IPT5b*/*LOG1b*) in stems, and that *SPL16/23* and *IPT5b/LOG1b* exhibit highly overlapping expression profiles in the phloem and cambium zones, we hypothesized that SPL16 and SPL23 might control CK levels to promote cambial activity in poplar. In support of this hypothesis, expression levels of *IPT5a, IPT5b,* and *LOG1b* were significantly downregulated in the stem bark of *spl16/23* mutants. Conversely, these genes were significantly upregulated in the stem bark of *CLE41b_pro_:SPL16* transgenic lines compared to WT plants (Figure 6A and B). Moreover, measurement of endogenous CK level revealed markedly lower tZ and iP CK contents in stems of *spl16/23* mutants than those in WT plants, while *CLE41b_pro_:SPL16* transgenic plants exhibited notably higher bioactive CK levels compared to WT (Figure 6C). These findings suggest that SPL16 and SPL23 act as positive regulators of CK biosynthesis in poplar stems.

**Figure 6.**
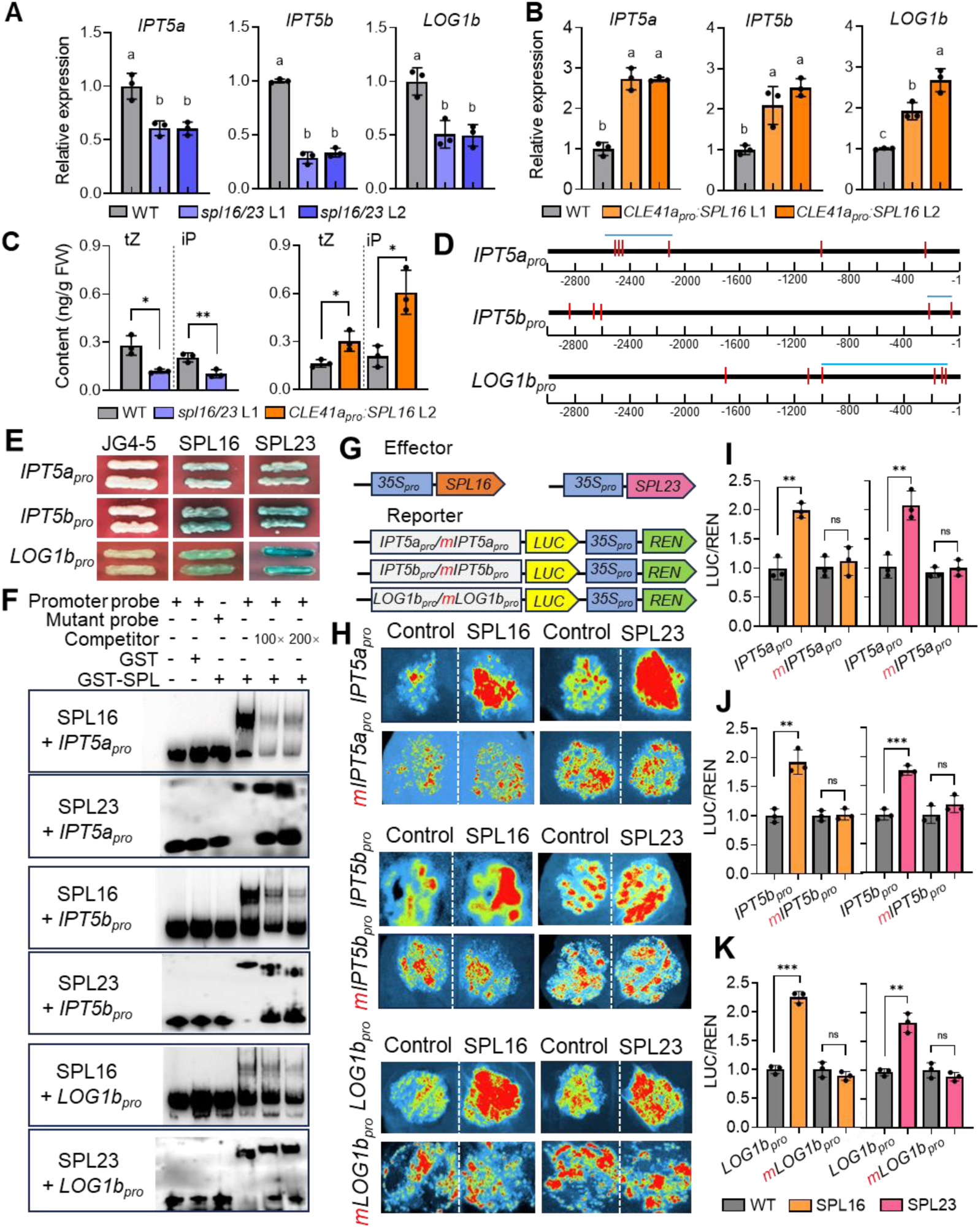
**SPL16 and SPL23 directly activate CK biosynthesis genes by binding to their promoters.** (**A**) Relative expression levels of *IPT5a*, *IPT5b* and *LOG1b* in the 7^th^ internodes of WT and *spl16/23* mutants. Data are mean ± SD from three different plants. (**B**) Relative expression levels of *IPT5a*, *IPT5b* and *LOG1b* in the 7^th^ stem internodes of WT and *CLE41b_pro_:SPL16* transgenic plants. Data are mean ± SD from three different plants. (**C**) Quantification of active CK (iP and tZ) contents in stem bark of the 7^th^ internodes from WT, *spl16/23* (L1) and *CLE41b_pro_:SPL16* (L2) transgenic plants. Data are mean ± SD from three different plants. (**D**) Identification of putative SPL binding sites (GTAC motifs) within 3kb upstream of the ATG start codon in the promoters of *IPT5a*, *IPT5b* and *LOG1b*. Horizontal blue lines denote the amplified promoter fragments of *IPT5a_pro_* (579 bp), *IPT5b_pro_* (183 bp) and *LOG1b_pro_* (926 bp), respectively, used in yeast one-hybrid (Y1H) assays. (**E**) Y1H results demonstrating the binding of SPL16 and SPL23 to the promoter fragment of *IPT5a*, *IPT5b*, and *LOG1b*. (**F**) EMSA showing direct binding of GST-SPL16 and GST-SPL23SBP recombinant proteins to the biotin-labeled probes of *IPT5a*, *IPT5b*, and *LOG1b* promoters (about 60 bp). In the mutant probes, GTAC was replaced by CCGG. (**G**) Schematic representation of effector and reporter constructs used for transient expression assays. *SPL16/SPL23* expressed under the control of the *35S* promoter was used as the effector. The firefly luciferase gene *LUC* driven by the *IPT5a_pro_* (2,605 bp), *IPT5b_pro_* (2,846 bp) and *LOG1b_pro_* (1,709 bp) promoters, respectively, was used as the reporter. *Renilla* luciferase gene *REN* driven by the *35S* promoter were used as the internal control. Red “m” indicates mutation of GTAC to GGAA. (**H**) LUC assays showing the activation of *IPT5a*, *IPT5b*, and *LOG1b* promoter activities by SPL16/SPL23 in *Nicotiana benthamiana* leaves. (**I-K**) Dual luciferase assays confirming SPL16/SPL23 activation on *IPT5a*, *IPT5b*, and *LOG1b* promoters. Rluc activity was used as internal control. Data are means ± SD from three biological replicates. Statistics significance: Asterisks denote significant differences by Student’s *t*-test (\**P* < 0.05; \*\**P* < 0.01; \*\*\**P* < 0.001; ns, not significant). Different letters indicate significant differences (*P* < 0.05) by one-way ANOVA and LSD test for pairwise comparisons.

To investigate whether SPL16/23 directly activate CK biosynthetic genes, we first analyzed the *cis*-elements (GTAC) in the promoter sequences (∼3 kb upstream of the start codon) of *IPT5a/IPT5b/LOG1b*. As depicted in Figure 6D, multiple SPL binding motifs (5′-GTAC-3′) are present in their promoter regions. Yeast one-hybrid assays showed that both SPL16 and SPL23 bound to the promoter fragments of *IPT5a/IPT5b/LOG1b* harboring SPL binding sites (Figure 6E). To further confirm this binding, we performed gel electrophoresis mobility shift assays (EMSA) using purified SPL16-GST and SPL23SBP-GST recombinant proteins. The results revealed that SPL16 and SPL23 fusion proteins specifically bound to biotin-labeled promoter fragments of *IPT5a/IPT5b/LOG1b* with intact GTAC motifs, but not to those with mutated motifs (Figure 6F). Next, we conducted transient expression assays in *Nicotiana benthamiana* leaves to determine whether SPL16/23 directly activate transcription of *IPT5a*, *IPT5b* and *LOG1b*. Co-injection of *Agrobacterium* strains harboring *35S_pro_:SPL16*/23 effector and *IPT5a_pro_/IPT5b_pro_/LOG1b_pro_*-driven *LUC* reporter plasmids resulted in significant activation of the LUC reporter gene expression from these promoters, but not from those with mutated GTAC motifs (Figure 6G and H). Dual luciferase assays (LUC/REN) confirmed that SPL16 and SPL23 enhanced the transcription of *IPT5a*, *IPT5b* and *LOG1b* (Figure 6I-K). These findings support that SPL16/23 positively regulate cambial activity through enhancing *IPT5s/LOG1b*-mediated active CK biosynthesis in poplar stems.

### Phloem-specific expression of *IPT5b* and *LOG1b* rescues the cambium division defects in *spl16/23* mutants

To verify that *IPT5b* and *LOG1b* act genetically downstream of *SPL16* and *SPL23* in regulating cambium activity, we introduced *CLE41b_pro_:IPT5b* and *CLE41b_pro_:LOG1b* constructs into the *spl16/23* mutant background, respectively (Supplemental Figure S16A and S17A). Phenotypic analysis of cross-sections from *spl16/23* mutants and *CLE41b_pro_:IPT5b>>spl16/23* transgenic lines under normal light conditions revealed that phloem-specific overexpression of *IPT5b* restored the number of cambial cells in *spl6/23* mutants (Supplemental Figure S16C and D). Additionally, elevated *IPT5b* expression resulted in a wider xylem and an increased number of xylem cell layers compared to *spl6/23* mutants (Supplemental Figure S16E and F). Notably, the *CLE41b_pro_:IPT5b>>spl16/23* (L3), with the highest *IPT5b* expression, exhibited the greatest increase in cambial and xylem cell layers. Similarly, the introduction of *LOG1b* driven by the *CLE41b* promoter into *spl16/23* background also restored the defective cambial cell layers and improved wood development under normal light conditions (Supplemental Figure S17C-F). Subsequently, the *CLE41b_pro_:IPT5b>>spl16/23* (L3) and *CLE41b_pro_:LOG1b>>spl16/23* (L1) were elected for propagation and further analysis under both WL and simulated shade conditions.

Cross-sectioning and microscopic observations of the 7^th^ internode of different genotypes revealed that phloem-specific overexpression of *IPT5b* or *LOG1b* in the *spl16/23* mutants not only rescued the cambium division defects, but also slightly increased cambial cell layers compared to WT plants under normal WL conditions (Figure 7A and B). Under shade conditions (R/FR of 0.77), *CLE41b_pro_:IPT5b>>spl16/23* and *CLE41b_pro_:LOGb>>spl16/23* transgenic poplars maintained higher cambial proliferation activity than both *spl16/23* and WT plants (Figure 7A and B). Accordingly, overexpression of *IPT5b* and *LOG1b* in the *spl16/23* mutants restored the defective xylem development to levels comparable to those in WT plants (Figure 7C and D). These findings provide genetic evidence that SPL16 and SPL23 regulate cambial activity by activating the CK biosynthetic genes *IPT5b* and *LOG1b*, implicating the SPL16/23-IPT5/LOG1-cytokinin pathway in shade-mediated inhibition of cambial activity in woody species.

**Figure 7.**
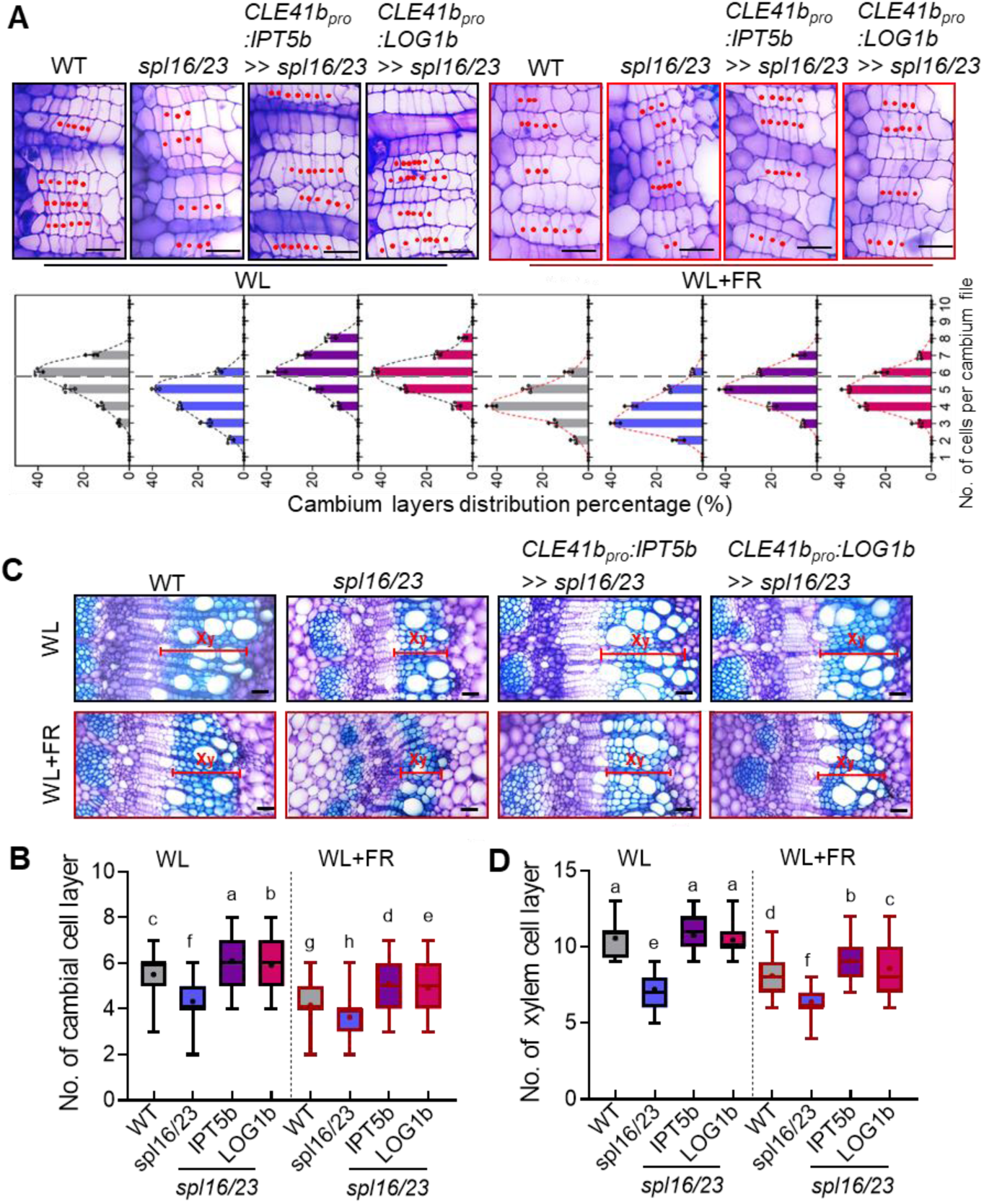
Phloem-specific expression of *IPT5b* or *LOG1b* rescues the vascular defects of *spl16/23* mutants. (**A**) Cambial phenotypes of *P. tomentosa* WT, *spl16/23* mutant, *CLE41b_pro_:IPT5b* >> *spl16/23*, and *CLE41b_pro_:LOG1b* >> *spl16/23* transgenic plants under white light (WL, R/FR = 10.67) and simulated shade (WL+FR, R/FR = 0.77) conditions (upper panel). The 7^th^ internodes of 10-week-old poplar plants of different genotypes were cross-sectioned, and stained by Toluidine blue. Red dots denote cambium cells. Bars = 50 μm. Frequency distribution of the number of cells per cambium cell file (lower panel). Values are mean ± SD from three different plants. (**B**) Quantification of the number of cambium cell layers in WT, *spl16/23* mutant, *CLE41b_pro_:IPT5b* >> *spl16/23*, and *CLE41b_pro_:LOG1b* >> *spl16/23* transgenic plants under both WL and WL+FR conditions. Values are mean ± SD (n = 100). (**C**) Xylem phenotypes in the 7^th^ internodes of WT, *spl16/23* mutant, *CLE41b_pro_:IPT5b* >> *spl16/23* and *CLE41b_pro_:LOG1b* >> *spl16/23* transgenic plants under both WL and WL+FR conditions. Xy, xylem. Bars, 50 μm. (**D**) Number of xylem cell layers in the 7^th^ internodes of different genotypes. Data are mean ± SD (n = 100). Whiskers indicate the minimum and maximum values, black lines within boxes denote the median values, and means are indicated by plus symbols. Different letters indicate significant differences (*P* < 0.05) by one-way ANOVA and LSD test for pairwise comparisons.

### Shade-induced miR156 expression contributed to the shade-induced inhibition of cambial activity

Considering that miR156 expression is modulated by light quantity and quality changes (Xie et al., 2017; Xu et al., 2021), and that *SPL16/SPL23* are regulated by miR156 at a post-transcriptional manner, we further tested whether simulated shade (low R/FR) could affect cambial activity through regulating miR156 transcript levels. We measured the abundance of mature miR156 in the stem bark of WT plants grown under normal WL and WL+FR (R/FR = 0.77) conditions using stem-loop qPCR. The results showed a significant increase in miR156 levels in the stems of shade-treated plants compared to the WL controls (Figure 8A). Consistent with the previous report that miR156 controls *SPL16/23* expression at the post-transcriptional level in poplar leaves (Wei et al., 2024), we observed downregulation of *SPL16* and *SPL23* in the stem bark of *MIR156a*-OE transgenic plants (L1, L2) compared to WT plants (Figure 8B). Consistently, expression levels of the downstream CK biosynthesis pathway genes *IPT5a*, *IPT5b*, and *LOG1b* were significantly decreased in these *MIR156a*-OE lines (Figure 8B). Furthermore, the levels of active CKs (iP and tZ) were much lower in the stem bark of *MIR156a-*OE stems compared to those in WT stems (Figure 8C).

**Figure 8.**
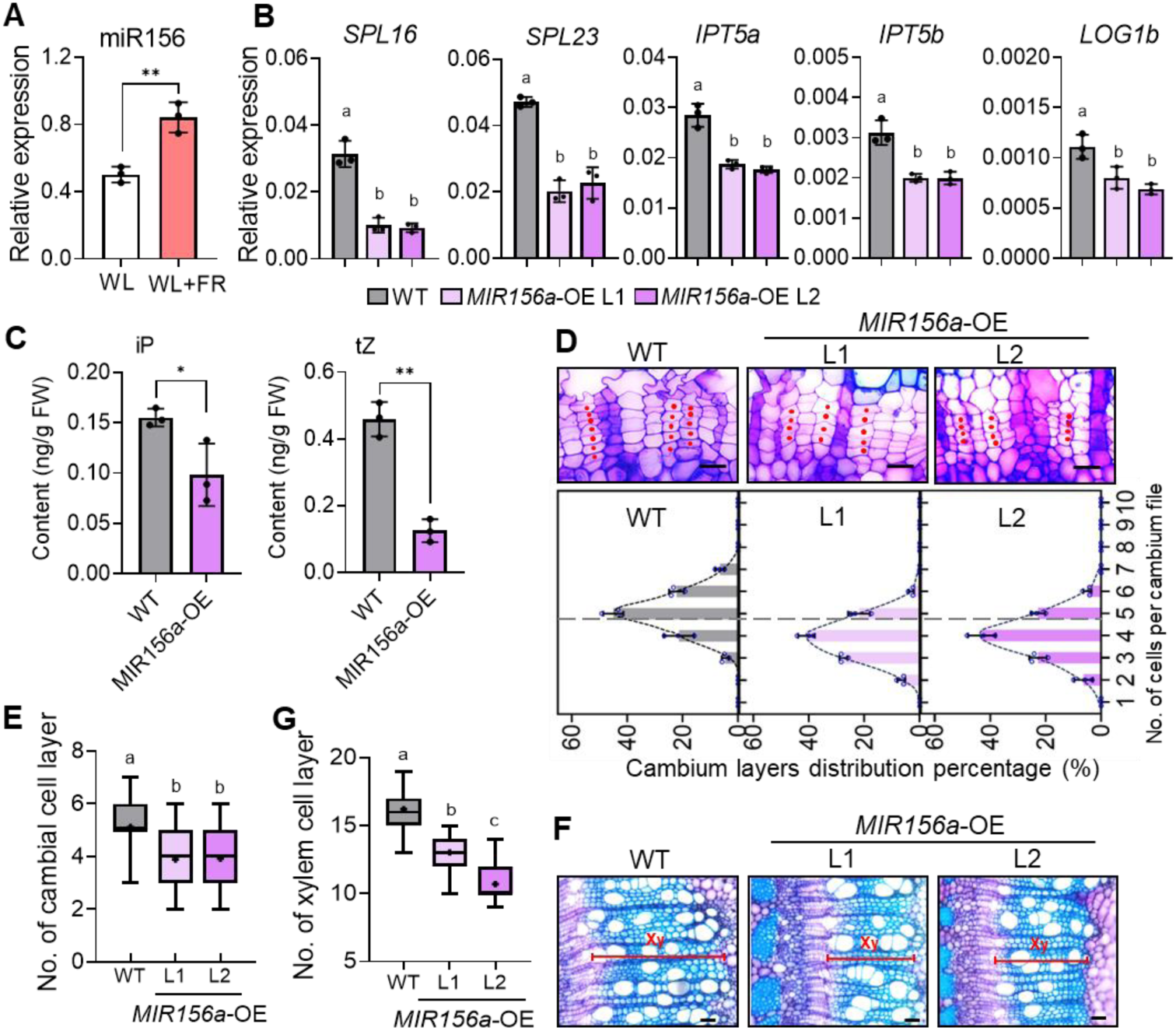
**Shade-induced miR156 expression contributed to the shade-induced inhibition of cambial activity.** (**A**) Abundance of mature miR156 in the 6^th^ internode of WT *P. tomentosa* plants under simulated shade (WL+FR). Data are mean ± SD from three different plants. (**B**) Expression levels of *SPL16* and *SPL23* and downstream CK biosynthesis genes *IPT5a*, *IPT5b*, and *LOG1b* in the 7^th^ internode of WT and *MIR156*-OE transgenic lines. Data are mean ± SD from three different plants. (**C**) Quantification of active CK (iP and tZ) contents in stem bark of the 7^th^ internodes from poplar WT and *MIR156a*-OE transgenic plants. Data are mean ± SD from three different plants. (**D**) Cambial phenotypes in cross-sections of the 7^th^ internodes of WT and *MIR156a*-OE transgenic lines under normal white light (WL) conditions (upper panel). Stem cross-sections were stained by Toluidine blue. Bars = 20 μm. Frequency distribution of the number of cells per cambium cell file (lower panel). Values are mean ± SD from three different plants. (**E**) Number of cells per cambium cell file of WT and *MIR156a*-OE transgenic plants. Values are mean ± SD (n = 120). (**F**) Xylem phenotypes in cross-sections of the 7^th^ internodes from WT and *MIR156a*-OE transgenic plants. Xy, xylem. Bars = 50 μm. (**G**) Number of xylem cell layers in the 7^th^ internode of WT and *MIR156a*-OE transgenic plants. Values are mean ± SD (n = 100). Whiskers indicate the minimum and maximum values, black lines within boxes denote the median values, and means are indicated by plus symbols. Statistics significance: Asterisks denote significant differences by Student’s *t*-test (\**P* < 0.05; \*\**P* < 0.01). Different letters indicate significant differences (*P* < 0.05) by one-way ANOVA and LSD test for pairwise comparisons.

These findings suggest that shade-mediated suppression of *SPL16/23* may also involve a miR156-dependent pathway.

To examine the effect of miR156 on cambial activity and xylem development, we compared the stem vascular phenotypes between WT and *MIR156*a-OE lines. Histological analysis of the cross-sections at the 7^th^ internode revealed that *MIR156a*-OE plants displayed impaired cambium proliferation, with a marked reduction in the number of cambial cells compared to WT plants (Figure 8D and E). Consequently, elevated expression of *MIR156a* led to a reduction in secondary xylem width and fewer cells per xylem cell file relative to the WT control (Figure 8F and G). These results support the notion that shade-induced miR156 levels in poplar stem partially contribute to the shade-inhibition of cambial activity and xylem differentiation during secondary growth.

## Discussion

### Shade inhibits cambial activity during secondary growth in *Populus* trees

Forest trees, particularly in high-density plantations, frequently face the challenge of shading by neighboring plants. While it has been well-documented that high stand density adversely impacts wood formation (Benomar et al., 2012), the molecular mechanisms for the shade-regulation of cambial activity and radial growth in trees remain largely unexplored. In this study, we demonstrated that *Populus* trees, upon sensing neighbor proximity signal (reduced R/FR ratio) in high stand density or when subjected to simulated shade treatment, prioritize internode elongation at the expense of radial growth. This leads to the suppression of cambial proliferation and secondary xylem development (Figure 1). Such an adaptive response likely enhances the competitive ability of trees in dense stands by reallocating resources towards vertical growth, thereby improving their access to light.

Extensive studies in Arabidopsis and cereal crops have elucidated the key roles of phytochrome photoreceptors, particularly phyB, and blue light photoreceptors (CRY1 and CRY2) in regulating shade avoidance responses (Liu et al., 2021). PIF transcription factors act as pivotal regulatory hubs downstream of these photoreceptors to integrate multiple environmental cues (e.g. light and temperature) and hormonal signals to trigger adaptive responses (Choi and Oh, 2016; Ballaré and Pierik, 2017). In *Populus* species, downregulation of *PHYB1/2* or knockout of *CRY1a/b* results in internode elongation, closely resembling the shade-induced phenotypes (Ding et al., 2021; Chen et al., 2024). Conversely, transgenic poplar plants overexpressing *PHYB1/PHYB2* exhibited shorter plant height and an attenuation of internode elongation under shade-mimicking conditions (Sun et al., 2024). *P. tomentosa* harbors eight PtoPIF genes, which exhibit both overlapping and distinct roles in controlling internode and leaf petiole elongation, as well as leaf angle adjustments in response to simulated shade (Sun et al. 2024). Notably, transgenic poplars overexpressing *PtoPIF3.1*/*PtoPIF3.2* displayed elongation of internodes and leaf petioles, and enhanced responses to shade through an auxin biosynthesis pathway (Sun et al. 2024). PtoPIF8.1 and PtoPIF8.2 seem not to mediate shade-induced elongation growth, rather play a role in shade-mediated inhibition of branching through indirectly promoting the expression of *BRANCHED1* (*BRC1*), a key negative regulator of bud outgrowth (Ding et al., 2021; Sun et al., 2024). Despite these insights, the specific roles of PtoPIFs in wood formation remain largely unexplored in poplar, highlighting an important area of future research.

In *Arabidopsis*, the phyB-PIF4 module has been reported to promote the elongation of inflorescence stems while simultaneously inhibit vascular development and secondary cell wall (SCW) formation under either high temperature or shade conditions (Luo et al., 2022; Wei et al., 2023). If the shade-inhibition of cambial activity in trees was mediated by phytochrome or cryptochrome signaling, one might expect that overexpression of photoreceptors or mutation of *PIFs* would stimulate cambial cell proliferation. Contrary to this expectation, overexpression of *Populus CRY1* suppressed stem radial growth by reducing the numbers of cambium cell layers and early developing xylem cells in mature stems, while the *cry1a/b* mutants displayed opposite phenotypes (Chen et al., 2024). Yet it was shown that the transition of primary to secondary growth was advanced in *CRY1a*-OE transgenic plants (Chen et al., 2024). Given the established roles of PtoPIF3.1/3.2 and PtoPIF8.1 in controlling shade avoidance responses, we evaluated their functions in cambial activity. Anatomical results showed that knocking out *PtoPIF3.1* or *PtoPIF3.2* impaired cambial proliferation and xylem development in poplar stems (Supplemental Figure S4), whereas mutation of *PtoPIF8.1* barely affected cambial activity and wood formation (Supplemental Figure S5). These findings implicate that PtoPIF3s and PtoPIF8.1 are unlikely mediators of the shade-inhibition of cambial activity in poplar, though the involvement of other PtoPIF transcription factors cannot be ruled out. Notably, distinct from Arabidopsis PIF4 and PIF5 that act as major regulators for cell elongation, overexpression of the *Populus* homolog *PIF4* strongly inhibited shoot growth, ultimately leading to lethality in poplar (Ding et al., 2021). Thus, further study is needed to elucidate the function and regulatory mechanism of PtoPIF4 in *Populus*, which may reveal the functional divergence of PIF4 homologs between herbaceous and woody species. Collectively, these results indicate that low R/FR ratio may regulate cambial activity and secondary growth by a mechanism that is different from the PIF-centered regulatory networks that govern typical SAS traits, such as stem elongation and suppressed branching.

### SPL16/23-mediated CK regulation underlies shade-induced inhibition of cambial activity

In annual plants, the miR156-SPL regulatory module plays pivotal roles in diverse plant growth and development processes, including vegetative and reproductive phase change, tillering/branching, leaf development, inflorescence architecture, and stress responses (Wang and Wang, 2015). In *Populus* trees, the miR156-SPL module serves as a conserved age-dependent switch determining the juvenile-to-adult phase change (Lawrence et al., 2021). Recent studies have provided critical insights into the roles of miR156-SPLs in photoperiodic control of growth cessation in poplar (Liao et al., 2023; Wei et al., 2024). However, the involvement of SPL transcription factors in wood formation has yet to be elucidated. Interestingly, increasing evidence suggests that the miR156-SPL module acts downstream of phytochrome-mediated shade signaling to regulate shade avoidance responses (Xie et al., 2017, 2020). Besides, transcript levels of miR156/157 and their *SPL* targets are regulated by low light (LL) intensity, delaying vegetative phase change (Xu et al., 2021). In this study, we screened *Populus SPLs* with high expression in woody tissues and examined their expression responses to simulated shade (low R/FR ratio). We focused on miR156-targeted *SPL16* and *SPL23* homologs, which are predominantly expressed in the phloem and cambium ray initials. Both *SPL16* and *SPL23* expression levels were significantly downregulated by simulated shade treatment (Figure 2). Consistent with this observation, the *Populus spl16* and *spl23* single and double mutants exhibited impaired cambial activity and reduced secondary xylem development (Figure 3A-D; Supplementary Figure S7). Conversely, *SPL16* overexpression in the phloem enhanced cambial proliferation and alleviated shade-induced inhibition of cambial activity (Figure 3E and F). These findings support that shade-induced suppression of *SPL16/23* expression levels in poplar stems are accountable for the shade-inhibition of cambial activity. It is also worth noting that *SPL20* and *SPL25*, which group with *SPL16/SPL23* in the *SPL3/5* clade, displayed reduced transcript levels under low R/FR conditions (Supplemental Figure S6A and C). This suggests potential functional redundancy or cooperation among these homologs. Indeed, *SPL16*, *SPL23*, *SPL20*, *SPL25*, and *SPL24* function cooperatively in controlling short photoperiod-induced growth cessation in *Populus* (Liao et al., 2023). Further investigation is needed to determine whether SPL20 and SPL25 also contribute to the regulation of cambial activity under canopy shade.

Cytokinin (CK) is a key regulator of various plant growth processes, including shoot regeneration, leaf development, stem secondary growth, and adaptive responses to altered environmental stresses (Kieber and Schaller, 2014; Nguyen et al., 2021). In poplar, CKs are enriched in the secondary phloem, where it regulates cambial activity in a noncell-autonomous manner (Immanen et al., 2016; Fu et al., 2021). The interaction of CK and light signaling has been documented. In *Arabidopsis*, canopy shade (low R/FR) causes an arrest in leaf development due to auxin-induced cytokinin inactivation mediated by the CK oxidase AtCKX6 in developing primordia (Carabelli et al., 2007). Analogous to the phenomena, our study found that simulated shade treatment inhibited leaf expansion in poplar trees. This inhibition was accompanied by a reduction in active CK (iP and tZ) levels in both leaves and stems, correlating with suppressed expression of CK biosynthesis genes *IPT5a*, *IPT5b*, and *LOG1b* (Figure 4, Supplemental Figure S10). We propose that shade-mediated reduction in CK levels across the whole plant contributes to the inhibition of leaf expansion and radial growth under canopy shade. Supporting this hypothesis, CRISPR/Cas9-mediated disruption of *IPT5a/IPT5b* led to smaller leaves and reduced cambial activity, while overexpression of *IPT5b* or *LOG1b* in the phloem of transgenic poplars enhanced cambial cell proliferation and counteracted shade-induced inhibition of cambial activity (Figure 4 and 5). These findings suggest that simulated shade modulates cambial activity in poplar by controlling CK levels in the stems, providing critical insights that simulated shade modulates cambial activity through controlling cytokinin concentration in poplar stems.

Importantly, our study establishes that SPL16 and SPL23 serve as a molecular connection between the shading signal response and the reduced cytokinin levels in poplar stems. The active CK contents were decreased in the *spl16/23* mutant stems, associated with decreased expression levels of the CK biosynthesis genes *IPT5a, IPT5b* and *LOG1b.* Molecular and biochemical studies corroborated that SPL16 and SPL23 directly bind to the promoters of *IPT5a*, *IPT5b,* and *LOG1b* to activate their expression, stimulating CK-mediated cambial activity (Figure 6). Notably, the defective cambial phenotypes of *spl16/23* mutants can be restored by the phloem-specific overexpression of *IPT5b/LOG1b* (Figure 7). Therefore, our study sheds light on the pivotal role of the SPL16/23-IPT5/LOG1-cytokinin pathway in shade-mediated suppression of cambial activity. In line with our findings, the regulatory relationship between SPL regulators and CK biosynthesis and signaling has been reported in several contexts. In apple, MdSPL14, forming a regulatory complex with MdWRKY24, directly represses the expression of *MdCKX5* and *MdARR6*, encoding a CK degradation enzyme and a type-A negative regulator of CK signaling pathway, respectively, to promote leaf development during vegetative phase transition (Jia et al., 2024). In rice, OsSPL14/IPA1 (IDEAL PLANT ARCHITECTURE 1) could directly regulate CK and auxin-related genes (*LOG* and *PIN1b*) to control plant architecture (Lu et al., 2013). Recent study also revealed that *OsSPL14/IPA1* was induced by exogenous CK treatment via type-B OsRR21-mediated transcriptional activation. IPA1 then directly activate the expression of CK receptor gene *OHK4*, forming a positive feedback circuit in regulating rice inflorescence architecture (Chun et al., 2023). Conversely, UNBRANCHED3 (UB3), a maize ortholog of OsSPL14, suppressed vegetative and reproductive branching by inhibiting CK biosynthesis and signaling (Du et al., 2017). Future research should explore whether SPL16 and SPL23 also regulate CK oxidases (CKX) and other CK signaling genes in poplar, and whether these transcription factors form a feedback regulatory loop with CK signaling in cambial activity regulation.

### Shade inhibits *SPL16/23* expression at both transcriptional and post-transcriptional levels

Accumulating evidence shows that miR156 levels are controlled by endogenous developmental signals and various environmental factors, such as light intensity, light quality (R/FR ratio), high temperatures, cold, and drought stress (Stief et al., 2014; Xie et al., 2017; Xu et al., 2021). In *Arabidopsis*, low light irradiance delays phase change by enhancing miR156 expression and reducing expression of miR156-targeted *SPL* through both miR156-dependnet and miR156-independent pathways (Xu et al., 2021). Unlike the effect of end-of-day FR light (EOD-FR) that represses miR156 expression and alleviates downstream *SPL* genes (Xie et al., 2017), our findings demonstrate that low R/FR ratio, a signal for canopy shade, increases miR156 expression and simultaneously reduces multiple miR156-regulated *SPL* genes in poplar stems (Figure 8A, Supplementary Figure S6C). The functional relevance of shade-induced miR156 expression on cambial activity is evidenced by the observations that transgenic poplars overexpressing *MIR156a* exhibited reduced CK levels and impaired cambial division and xylem development (Figure 6). This suggests that canopy shade may extends the juvenile phase and inhibits secondary growth in densely planted trees, prioritizing vertical growth to compete more effectively for light under dense canopy. In line with this speculation, it has been shown in poplar that juvenile leaves are photosynthetically more efficient than adult leaves under low light irradiance (Lawrence et al., 2020). It is also worth noting that juvenile poplar seedlings or transgenic poplars overexpressing *MIR156* exhibited prolonged vegetative growth under growth-restrictive short days, thus allowing better climate acclimation (Liao et al., 2023). These findings highlight the pivotal roles of the miR156-SPL module in integrating multiple stress responses with development. Additionally, our study shows that simulated shade significantly suppresses GUS expression driven by the *SPL16* and *SPL23* promoters (Figure 2D and E), providing strong evidence that low R/FR ratios suppress *SPL16/23* expression at both transcriptional and post-transcriptional levels. Further analyses will be carried out to uncover what upstream signaling pathways coordinately regulate the expression of *MIR156* and *SPLs*.

Given that low light negatively affects photosynthesis, while exogenous sugar application could partially restore the delay in vegetative phase transition (Xu et al., 2021). This aligns with earlier findings that glucose or sucrose treatment reduces the abundance of the endogenous *pri-MIR156A* and *pri-MIR156C* transcripts in *Arabidopsis* (Yu et al., 2013; Yang et al., 2013). Additionally, *SPL9* transcription is directly activated by PAP1/MYB75, whose expression is specifically induced by sucrose, suggesting a miR156-independent pathway in sugar-mediated vegetative phase transition (Meng et al., 2021). Interestingly, photosynthesis-derived sugars and exogenous sucrose treatment can stimulate de novo cytokinin biosynthesis, linking sugar signaling to CK metabolism (Kiba et al., 2019; Salam et al., 2021), implicating that the miR156-SPL regulatory module may bridge sugar singling toward cytokinin metabolism. Supporting this notion, shade-induced reductions in light intensity have been shown to deplete nonstructural carbohydrate reserves, primarily starch, in the stems of *P. nigra* trees (Tomasella et al., 2021). These observations suggest that shade-mediated changes in sugar levels could influence the expression of miR156-SPLs, thereby regulating CK levels and cambial proliferation activity. However, the impact of simulated shade on sucrose and glucose levels in the phloem and cambium zone, as well as their influence on the transcription of MIR156 and *SPL16/SPL23* genes, remains to be elucidated.

Notably, trees in boreal and temperate regions exhibit distinct seasonal patterns in cambial activity and wood formation, largely associated with fluctuations in photosynthetic sugar production. These seasonal variations suggest that cambial division and wood formation may be closely linked to sucrose concentrations in the stems (Uggla et al, 2001). Our re-analysis of transcriptome libraries from *Populus* during different cambial stages (dormant, reactivating, and active) reveals that miR156 levels peak during winter dormancy, while *SPL16/23/25/17/15* transcripts are more abundant in active cambial zones (Chen et al., 2021). This pattern suggests that the miR156-SPL module may play a critical role in regulating seasonal cambial growth and wood formation in perennial woody trees. Future studies should explore whether the miR156-SPL module acts as a crucial link between sugar sensing and downstream pathways, particularly CK biosynthesis, to modulate cambial activity in response to seasonal changes. This line of inquiry could provide valuable insights into the molecular mechanisms underlying the adaptation of cambial growth to seasonal fluctuations in sugar availability in trees.

Based on our results, we propose a schematic model for the shade-mediated inhibition of cambial activity during secondary growth in poplar stems (Figure 9). *SPL16* and *SPL23*, which are localized in the phloem, play a crucial role in maintaining CK levels by directly activating CK biosynthesis genes (*IPT5a*, *IPT5b*, *LOG1b*) under normal light conditions (high R/FR ratios). This activation supports CK’s role in promoting cambial proliferation and secondary growth. In contrast, under high stand density or simulated shade conditions (low R/FR ratios), which signal neighbor proximity, the expression of *SPL16* and *SPL23* in the phloem and cambium region is repressed both transcriptionally and post-transcriptionally, primarily through miR156. This repression leads to the downregulation of *IPT5a*, *IPT5b*, and *LOG1b*, culminating in decreased CK levels in the secondary phloem and cambium zone. Consequently, this reduction in CK levels inhibits cambial activity and impairs wood formation in poplar stems. These findings elucidate the mechanistic basis of how shade suppresses cambial activity and radial growth in woody species, offering insights for future molecular breeding strategies aimed at developing shade-tolerant trees.

**Figure 9.**
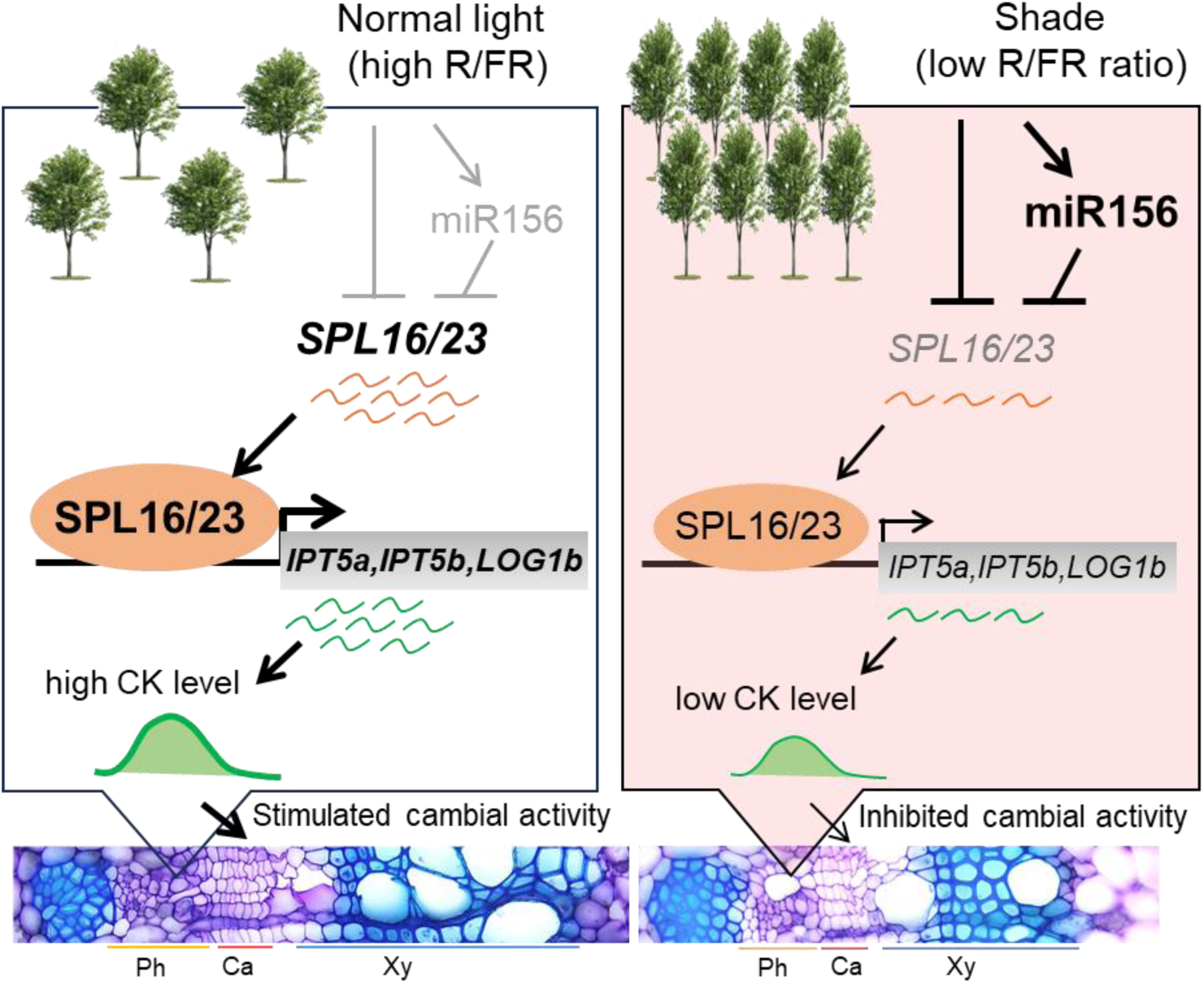
**Schematic model of simulated shade inhibits cambial activity in poplar stems via suppressing the SPL16/23-mediated cytokinin pathway.** This model illustrates the role of SPL16 and SPL23 as key regulators of cambial activity in *Populus* by maintaining cytokinin (CK) levels. Under conditions of low stand density or normal light (high R/FR ratio), *SPL16* and *SPL23*, localized predominantly in the phloem, play a crucial role in maintaining CK levels by directly activating CK biosynthesis genes (*IPT5a*, *IPT5b*, and *LOG1b*), and in turn promoting cambial proliferation and secondary xylem development in *Populus* stems. In contrast, under high stand density or simulated shade conditions (low R/FR ratios), the expression of *SPL16* and *SPL23* is repressed both transcriptionally and post-transcriptionally through miR156. This repression leads to the downregulation of *IPT5a*/*IPT5b*/*LOG1b* and ultimately decreased CK levels in the phloem and cambium zones. Consequently, this reduction in CK levels inhibits cambial activity and impairs wood formation in poplar stems. Arrows indicate activation; while bars indicate repression. Ca, cambium; Ph, phloem; Xy, xylem.

## Materials and methods

### Plant materials and growth conditions

*Populus tomentosa* Carr. (Clone 741) plants were micropropagated in vitro on Woody Plant Medium (WPM, Phyto Tech). The transgenic poplar lines, including the *spl16* single mutant, *spl16/23* double mutants, and *MIR156a*-OE, were previously described by Wei et al. (2024). The *spl23, pif8.1* and *ipt5a/b* mutants were generated in this study. Genotyping was performed by amplifying sequences harboring the CRISPR target sites using genomic DNA (gDNA) extracted from the corresponding mutants, followed by sequencing to confirm the mutations. Both wild-type (WT) and transgenic poplar plants were simultaneously propagated. After one month of rooting in tissue culture bottles, the plantlets were transferred to plastic pots (5 cm x 5 cm) filled with organic substrates. Following a three-week acclimatization period, the plants were transplanted into larger plastic pots with the same organic substrate and grown in a greenhouse under long-day conditions (16 h light/8 h dark, 24°C, 60% humidity).

### Simulated shade treatments

Poplar plants were cultivated under normal light conditions, consisting of six white lights (6WL, 3629 lux, R/FR ratio of 10.69). For simulated shade treatments, 10-week-old poplar plants were either maintained under normal light (6WL) as a control or subjected to low R/FR ratios by supplementing the light with far-red (FR) lights. The treatments included: (1) 6WL+2FR, R/FR ratio = 0.77; (2) 6WL+3FR, R/FR ratio = 0.46; (3) 4WL+2FR, R/FR ratio = 0.55. The simulated shade treatments were conducted for 8 to 14 days, as specified in the figure legends. Light intensity and quality were measured using an HR-550 spectrometer (HiPoint Inc., www.hipoint.com.tw).

### High stand density trials in field conditions

*P. tomentosa* WT plants were grown in plastic pots under low stand density (LSD) and high stand density (HSD) in filed conditions (Chongqing, China), starting in May, 2022. For LSD (16,448 lux, R/FR ratio of 1.00), two pots were spaced apart (35 cm × 35 cm) on a tray. For HSD (12,868 lux, R/FR ratio of 0.62), four pots were placed closely together (17 cm × 17 cm). Larger poplar plants were positioned around the trays to ensure canopy shade. Light intensity and quality were measured between neighboring plants at 16:00 on a sunny day using an HR-550 spectrometer. After four weeks, plants were photographed and subjected to phenotypic analysis.

### Plasmid construction

CRISPR mutants in poplar were generated by designing sgRNA targets within the first and second exons of the target genes. The CRISPR/Cas9 target sites were amplified from *P. tomentosa* genomic DNA, confirmed by sequencing, and cloned into the pYLCRISPR/Cas9 vector (Ma et al., 2015). For phloem-specific overexpression, the coding sequences (CDS) of *SPL16*, *IPT5b*, and *LOG1b* were PCR-amplified from *P. tomentosa* cDNA and inserted downstream of the *CLE41b* promoter using homologous recombination (ClonExpress II MultiS One Step Cloning Kit, Vazyme). Promoter:GUS reporter lines were generated by amplifying the promoter sequences of *SPL16_pro_* (2,192 bp), *SPL23_pro_*(2,185 bp), *IPT5b_pro_* (2,797 bp), and *LOG1b_pro_*(1,709 bp) from *P. tomentosa* gDNA and cloning them into the pCXGUS-P vector (Chen et al., 2009). For recombinant protein production, the coding sequences for SPL16 full-length protein and the coding sequences for SPL23 SBP domain were fused with GST in the *pGEX-4T-1* vector at the *EcoR*I restriction site to generate the *GST-SPL16* and *GST-SPL23SBP* construct. All the primers used for the constructs above are shown in Supplemental Data Set S1.

### Plant transformation and genotyping

*P. tomentosa* was genetically transformed via Agrobacterium-mediated infiltration of leaf disks as described previously (Jia et al., 2010). Notably, the *CLE41b_pro_:IPT5a* and *CLE41b_pro_:LOG1b* constructs were transformed into both WT and *spl16/23* mutant background. For genotyping of CRISPR mutants, DNA fragments harboring the target sequences were amplified by gene-specific primers from the gDNA of putative mutants, and then the PCR products were sequenced to identify homozygous knockout mutants. The expression levels of *SPL16*, *IPT5b* and *LOG1b* in stems of the corresponding overexpressing plants were detected by quantitative PCR (qPCR). Then the selected transgenic poplar plants were propagated together with WT for further phenotypic analysis.

### Cross-sectioning and histological staining

The 7^th^ internode (counted from the top) of *P. tomentosa* WT and transgenic poplar plants, unless otherwise indicated in the figure legend, were sampled for cross-sectioning with a vibrating blade microtome (VT1000s; Leica, Wezlar, Germany). The internodes were cut into 80 μm sections, and then were stained with 0.05% (w/v) toluidine blue for 5 min, followed by observation and photographing under a microscope (BX53; Olympus, Tokyo, Japan). The number of cambial cells, seen as undifferentiated flat cells, were counted from over 100 cambial cell files. The calculated frequency of anticlinal division represents the percentage of the number of anticlinal divisions in relation to the total cell number of the corresponding cambial cell file, according to the previously described method (Schrader et al., 2004; Nilsson et al., 2008). The number of xylem cells in each xylem cell file was counted from over 100 xylem cell files.

### GUS staining and MUG assays

Several transgenic lines of *P. tomentosa SPL16_pro_:GUS*, *SPL23_pro_:GUS*, *IPT5b_pro_:GUS*, and *LOG1b_pro_:GUS* reporter plants were identified by GUS histochemical staining, transgenic lines with strong GUS signal were propagated for further analysis. To examine how simulated shade affect promoter activities, *SPL16_pro_:GUS*, *SPL23_pro_:GUS*, *IPT5b_pro_:GUS*, and *LOG1b_pro_:GUS* reporter plants were grown under normal light (6WL, R/FR of 10.69) for 8 weeks, then were either maintained this light condition as controls or transferred to simulated shade (6WL+2FR, R/FR ratio of 0.77) for eight days. Then cross-sections of the 7^th^ internode from both WL and shade-treated plants were soaked in GUS staining solution. The stained cross-sections were washed three times with anhydrous ethanol: glacial acetic acid (3:1) at 65℃ to remove chlorophyll, followed by microscopic observation using an Olympus BX53 microscope. For quantitative quantification of GUS activity, MUG (4-methylumbelliferyl-beta-D-glucuronide) activity was determined using a fluorescence spectrophotometer (F-7000, Hitachi, Tokyo, Japan) according to the previously described method (Jefferson, 1987). Protein concentration was determined using the Bradford method (Kruger, 1996).

### Hormone quantification

Leaf samples and stem bark tissues from the 10-week-old WT poplar plants in normal light (R/FR of 10.67) and simulated shade (R/FR of 0.77) conditions were collected for measurement of auxin and cytokinin content. The stem bark was stripped off from the 6^th^ and 7^th^ internodes of WT (WL versus WL+FR) and the indicated transgenic poplar plants. Three different plants were analyzed, each sample containing approximately 0.3 g of stem bark tissues that were collected and immediately frozen in liquid nitrogen. Auxin and active cytokinins (iP and tZ) were detected based on the SCIEX Triple Quad™ 6500+ LC-MS/MS, platform (AB SCIEX, USA). Briefly, samples were ground to powder and extracted with 1 mL methanol/water/formic acid (15:4:1, v/v/v). The combined extracts containing added internal standards were evaporated to dryness under nitrogen gas stream using a concentrator (DC150-2, YouNing Instrument, CN). The samples were redissolved in methanol with 0.1% (v/v) formic acid, and filtered through a 0.22 μm filter for further HPLC-MS/MS analysis. The standards used were purchased from Olchemim Ltd. (Olomouc, Czech Republic) and TRC (Toronto, Canada). Data acquisition was performed using Analyst 1.6.3 software (AB Sciex). All metabolites were quantified using Multiquant 3.0.3 software (AB Sciex).

### RNA extraction and qPCR analysis

For screening the expression changes of *Populus SPL* family genes and other genes of interest in the phloem and cambium zone, the stem bark tissues were stripped off from the 5^th_^7^th^ internodes of 10-week-old *P. tomentosa* WT plants grown under normal light (6WL) and simulated shade (6WL+2FR, 6WL+3FR) treatment for 8 days. Three independent poplar plants were used for sampling. For identifying overexpression plants driven by the phloem-specific promoter, stem bark tissues were collected from the 7^th^ internodes of poplar WT and transgenic lines. Samples were immediately frozen in liquid nitrogen and stored at −80°C until use. Total RNA was extracted using Biospin Plant Total RNA Extraction Kit (Bioflux). First-strand cDNA was synthesized using Hifair AdvanceFast 1st Strand cDNA Synthesis Kit (Yeasen, Shanghai, China). qPCR was performed using Hifair Advanced One Step RT-qPCR SYBR Green Kit (Yeasen, Shanghai, China) according to the manufacturer’s instructions in a qTOWER3G IVD Real-Time PCR machine (Analytik Jena AG). Gene expression level was normalized to that of the reference gene *UBIQUITIN* (*UBQ*) using the 2^−ΔCt^ or 2^−ΔΔCt^ quantification method. The primers used are listed in Supplemental Data Set S1.

### Mature miRNA detection

The 6^th^ internodes of *P. tomentosa* WT plants under WL and WL+FR conditions were collected for total RNA extraction using TRIzol reagent (Invitrogen) according to manufacturer’s instructions. For detecting transcript levels of mature miR156, stem-loop qRT-PCR was performed as described previously (Kramer, 2011). U6 small nuclear RNA was used as an internal control for miR156 expression detection. Relative expression levels were calculated by the 2^-ΔCt^ method. The primers used are listed in Supplemental Data Set S1.

### RNA *in situ* hybridization

The sense and anti-sense fragments of *SPL16* (267 bp), *SPL23* (218 bp), *IPT5b* (154 bp), and *LOG1b* (196 bp) were amplified from *P. tomentosa* cDNA and labeled using a DIG RNA Labeling Kit (Roche) as probes. The primers used are listed in Supplemental Data Set S1. The 7^th^ internode of 2-month-old wild-type *P. tomentosa* plants was sampled for cross-sectioning. Stem cross-sections were then hybridized with the gene-specific sense and anti-sense probes of *SPL16*, *SPL23*, *IPT5b*, and *LOG1b*, respectively. *In situ* hybridization was performed according to the previously described method (Sang et al., 2012).

### Y1H

For yeast one-hybrid assay, the promoter fragments of *IPT5a_pro_*(579 bp), *IPT5b_pro_* (183 bp), and *LOG1b_pro_* (926 bp) were individually amplified from gDNA of *P. tomentosa* and ligated into *pLacZi2μ* vector to generate the constructs *IPT5a_pro_-2μ*, *IPT5b_pro_-2μ* and *LOG1b_pro_-2μ*, respectively. The JG-SPL16 and JG-SPL23 plasmids were previously reported (Wei et al., 2024). To test the binding of SPL16 and SPL23 to the promoters of *IPT5a, IPT5b* and *LOG1b* genes in yeast, combinations of the indicated pJG4-5 and pLacZi2μ plasmids were co-transformed into the yeast strain EGY48 and grown on the SD/-Ura/-Trp medium. Positive transformants were further streaked on the selection medium containing raffinose, galactose, and X-Gal (5-Bromo-4-Chloro-3-Indolyl β-D-Galactopyranoside) (Amresco, USA) for blue color development.

### Recombinant protein production

For recombinant protein production, the coding sequences for *SPL16* full-length protein and the coding sequences for SPL23 SBP domain were fused with GST in the *pGEX-4T-1* vector at the *Eco*RI restriction site to generate the *GST-SPL16* and *GST-SPL23SBP* construct. The *GST-SPL16* and *GST-SPL23SBP* constructs were individually transformed into the *Escherichia coli* strain Transette (TransGen Biotech, China). Then positive colonies were incubated in Luria-Bertain (LB) medium at 37 °C overnight and diluted at 1:100 with fresh LB medium and incubated at 37 °C for 3 h with OD_600_ of 0.6-0.8. After induction by 0.4 mM isopropyl β-D-thiogalactopyranoside (IPTG) at 37 °C, the cells were harvested by centrifugation at 4,000 × g for 10 min at 4 °C. For purification of *GST-SPL16* and *GST-SPL23SBP* proteins, the pellets were resuspended in PBS buffer (137 mM NaCl, 2.7 mM KCl, 10 mM Na_2_HPO_4_, 1.76 mM KH_2_PO_4_, pH 7.4). After being ultrasonicated, the supernatant of cell debris was recovered by centrifugation at 13,000 g for 60 min and then were purified using Glutathione-Sepharose resin (GE Healthcare, USA) according to the manufacturer’s protocol. The GST-SPL16 and GST-SPL23SBP recombinant proteins in elution buffer (20 mM reduced glutathione, 100mM Tris-Cl, pH 8.0) were dialyzed against the dialysis buffer (5 mM Tris-Cl, 50 mM KCl, pH 8.0), and were used for gel shift assay or stored at -80℃.

### EMSA

To examine the binding of SPL16 and SPL23 to the target promoters, two complementary oligonucleotides of *IPT5a/IPT5b/LOG1b* promoters (about 60 bp length) containing the SPL binding sites or mutated binding sequences (GTAC replaced by CCGG) were synthesized and labeled with biotin separately, and double strand probes were obtained by annealing the two oligonucleotides. The recombinant GST-SPL16 or GST-SPL23SBP protein was incubated with labeled probes at room temperature for 20 min, and the unlabeled probes were used as a competitor. The GST proteins were used as a negative control. The protein-DNA complexes were separated by electrophoresis on a native 4% acrylamide gel. The DNA was electroblotted onto nitrocellulose membranes and detected using a Chemiluminescent EMSA Kit (Beyotime, China). EMSA images were photographed using the Biostep Celvin S420 system (Biostep, Germany).

### Luciferase activity assays

For luciferase activity assays, the promoter sequences of *IPT5a_pro_* (2,605 bp), *IPT5b_pro_*(2,846 bp), and *LOG1b_pro_* (1,709 bp) were separately fused with the luciferase (*LUC*) gene in the *pGreenII0800* vectors (Biovector, USA). *Agrobacterium* harboring the *SPL16/23* expression plasmids and *IPT5a_pro_/IPT5b_pro_/LOG1b_pro_:LUC* plasmids were co-injected into *N. benthamiana* leaves. The SPL binding sites (GTAC) in these promoters were mutated as GGAA to generate *mIPT5a_pro_/mIPT5b_pro_/mLOG1b_pro_:LUC* plasmids. After incubating at 25 °C for 2-3 days, the injected leaves were detached and sprayed with 1 mM D-luciferin potassium salt solution (YEASEN, China). The luciferase luminescence was observed using a Fusion SL4 spectral imaging system (Vilber, Beijing, China). The luciferase activity was measured using the Dual Luciferase Assay kit (YEASEN, China) following the manufacturer’s protocol and the relative firefly luciferase activity was counted as the ratio of firefly to *Renilla* luciferase activity (LUC/REN) for each sample.

### Phylogenetic analysis

The amino acid sequences for SPL transcription factors, IPT and LOG proteins from *Arabidopsis thaliana* and *P. trichocarpa* were retrieved from Phytozome (https://phytozome.jgi.doe.gov/). An unrooted phylogenetic tree was constructed using MEGA9.0 with the statistical method of neighbor-joining method with bootstrap (1000 replicates) in MEGA9.0.

### Statistical analyses

The figures were produced using GraphPad Prism software (version 7.04). One-way ANOVA and the Least Significance Difference (LSD) test was adopted to analyze the significant differences (*P* <0.05) among multiple groups using the SPSS software (IBM SPSS Statistics 20). Student′s *t* test was adpoted to analyze the significant difference (\**P* <0.05; \*\**P* <0.01; \*\*\**P* <0.001) between two groups of data using the GraphPad Prism software. Statistical data are summarized in Supplemental Data Set S2.

## Accession numbers

Sequence data can be found in Phytozome Database under the following accession numbers: *SPL16* (Potri.011G055900), *SPL23* (Potri.004G046700), *IPT5a* (Potri.008G202200), *IPT5b* (Potri.004G046700), *LOG1b* (Potri.004G212200).

## Acknowledgments

This work was supported by the National Natural Science Foundation of China (32371903, 32101483), National Key Research and Development Program of China (2022YFD1201600), and Chongqing Major Special Project for Promoting Forestry through Science and Technology (2D-2022-2).

## Conflict of interest

No conflict of interest declared.

## Author contributions

H.W. and K.L. conceived and designed the experiments. X.X., J.D. and Y.L. performed the experiments. M.L., X.X., J.D. and Y.L collected the phenotypic data.

C.Z. and J.X. analyzed the data. H.W. and K.L. wrote the paper.

## Supplemental Figures

**Supplemental Figure S1.**
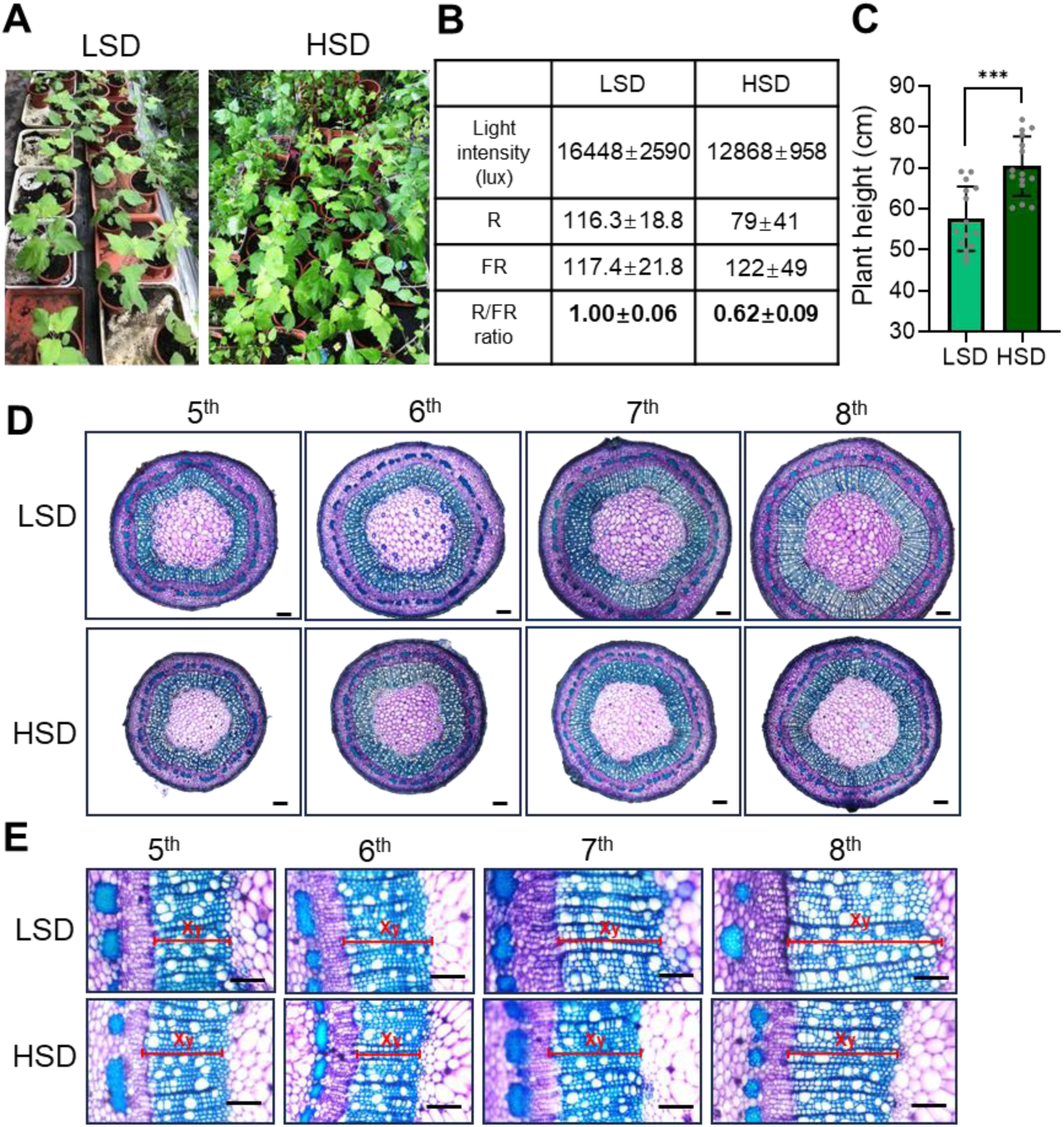
High stand density inhibits stem radial growth in *Populus* trees. (**A**) Experimental setups for low stand density (LSD) and high stand density (HSD) conditions. *Populus tomentosa* wild-type (WT) plants were grown in plastic pots on trays maintaining a distance between neighboring plants (35 cm x 35 cm) in LSD. In HSD, poplars were placed adjacent to each other on trays (17 cm x 17 cm) with larger plants around to provide canopy shade. Plant spacing was adjusted during growth. (**B**) Light intensity measurements of total light (lux), red (R), and far-red (FR) light between neighboring plants under LSD and HSD conditions, using an HR-550 spectrometer. The R/FR ratio is calculated to indicate shade signaling. (**C**) Plant height measurements of WT plants after two weeks of HSD treatment. Asterisks indicate significant differences (\*\*\**P* < 0.001) by Student’s *t*-test. (**D**) Cross sections of the 5^th^ to 8^th^ internodes of 2-month-old WT poplar plants grown in LSD and HSD conditions. Bars, 100 μm. (**E**) Close-ups of the xylem phenotypes of stem cross-sections shown in (**D**). Bars, 100 μm.

**Supplemental Figure S2.**
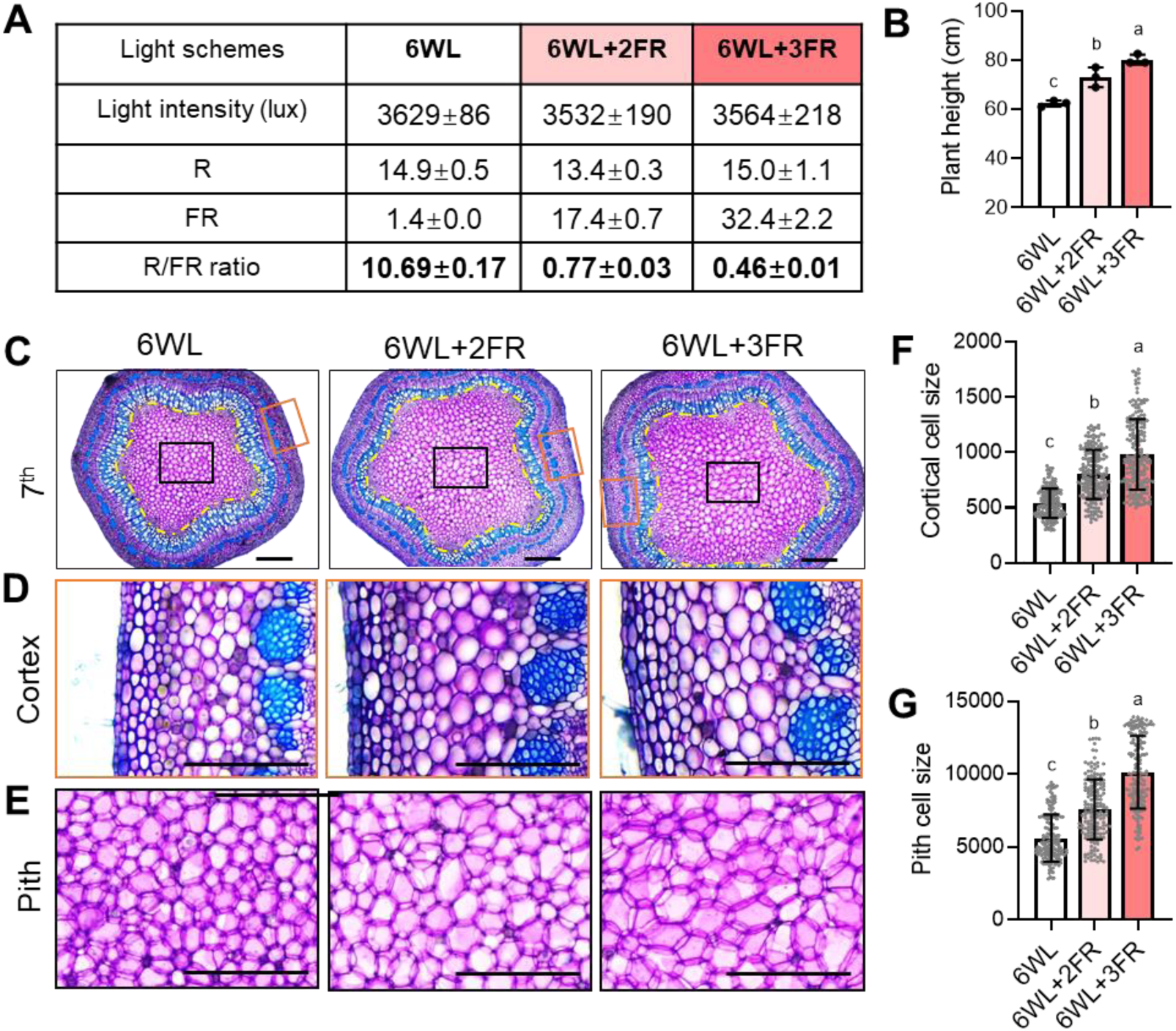
Simulated shade stimulates expansion of cortical and pith cells in stems of *Populus* trees. (**A**) Experimental setup for simulated shade treatment. *P. tomentosa* WT plants were initially grown in a greenhouse under normal white light conditions (6WL, R/FR ratio of 10.69) for 10 weeks. Poplar plants were then maintained under normal light (6WL) as controls or transferred to conditions with added far-red (FR) lights: 6WL+2FR (R/FR ratio of 0.77) or 6WL+3FR (R/FR of 0.46) for 8 days. Light intensity (lux), red (R) and far-red (FR) light intensities were measured using an HR-550 spectrometer. (**B**) Plant height measurements of of approximately 11-week-old WT poplars grown under normal light (6WL) and simulated shade conditions (6WL+2FR and 6WL+3FR). (**C**) Cross-sections of the 7^th^ internodes of WT poplars grown under normal light (6WL) and simulated shade conditions (6WL+2FR and 6WL+3FR). Bars, 100 μm. (**D**) Close-up images of cortical cells from the cross sections shown in (C), indicated by the orange rectangle. Bars, 50 μm. (**E**) Close-up images of pith cells from the cross sections shown in (C), indicated by the black rectangle. Bars, 50 μm. (**F**) Quantification of the cortical cell size in the 7^th^ internodes of WT poplars grown under normal light (6WL) and simulated shade conditions (6WL+2FR and 6WL+3FR) (n = 200). (**G**) Quantification of the pith cell size in the 7^th^ internodes of WT poplars grown under normal light (6WL) and simulated shade conditions (6WL+2FR and 6WL+3FR) (n = 170). Different letters indicate significant differences (*P* < 0.05) by one-way ANOVA and LSD test for pairwise comparisons.

**Supplemental Figure S3.**
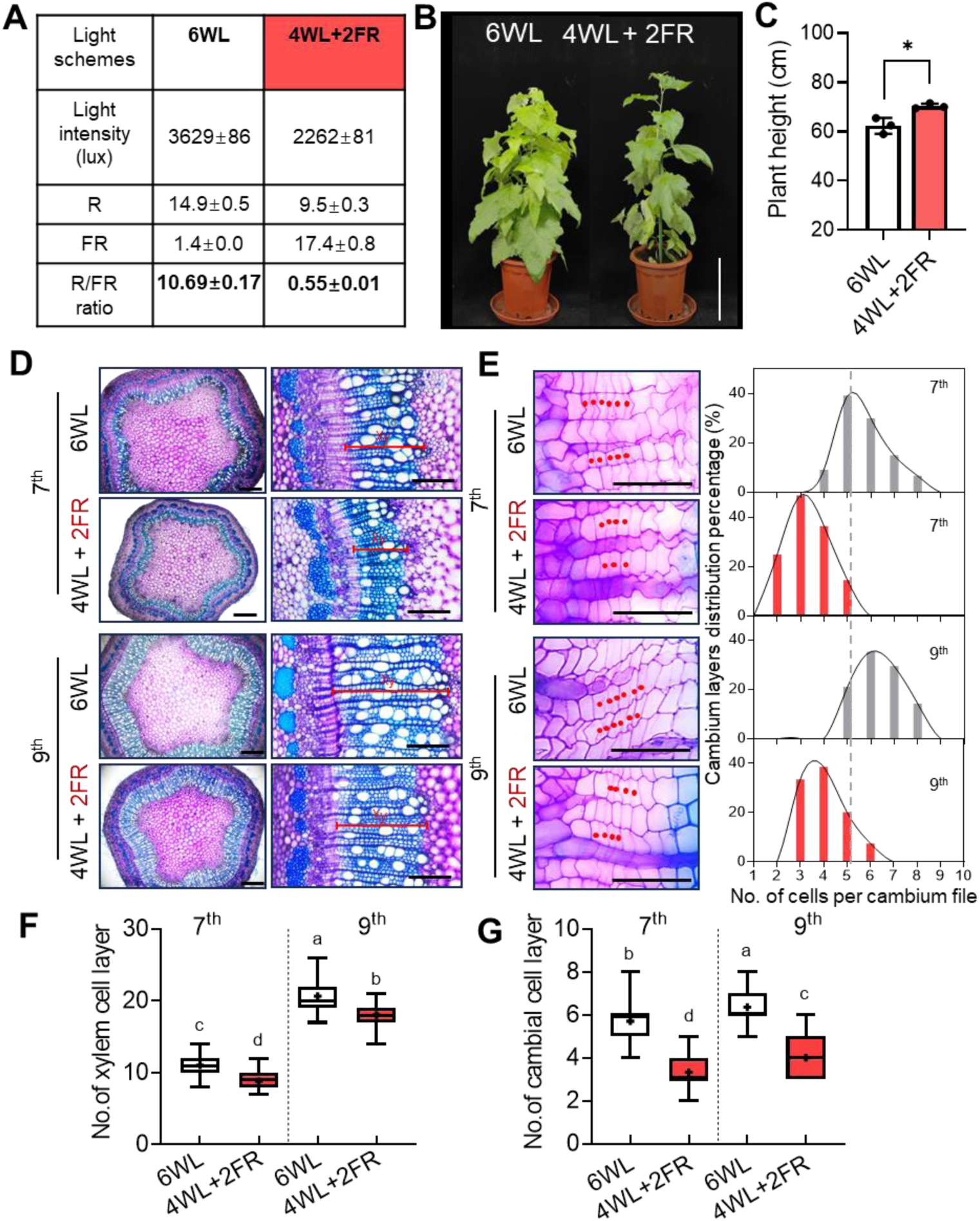
Decreased light intensity inhibits stem radial growth in *Populus* trees. (A) Experimental setup for simulated shade with reduced light intensity. Compared to the normal light condition (6WL, R/FR ratio of 10.69), conditions with 4 WL supplemented with 2 far-red lights (4WL+2FR, R/FR ratio of 0.55) was used to mimic high stand density conditions with lower light intensity and higher R/FR ratio. Light intensity (lux), red (R) and far-red (FR) light intensity were measured using an HR-550 spectrometer. (**B**) 10-week-old WT poplar plants were maintained under normal light (6WL) as controls or treated with 4WL+2FR for 8 days. Bar, 22 cm. (**C**) Measurements of plant height for WT poplar plans under 6WL and 4WL+FR conditions. Data are mean ± SD (n = 3), Asterisks denote significant differences by Student’s *t*-test (\*\*\**P* < 0.001). (**D**) Left panel: Toluidine blue-stained cross-sections of the 7^th^ and 9^th^ internodes of WT poplar plants under 6WL and 4WL+FR conditions. Bars, 500 μm. Right panel: Xylem phenotypes of WT poplar plants under 6WL and 4WL+FR conditions. Bars, 200 μm. (**E**) Left panel: Close-ups of cambial cells in the 7^th^ and 9^th^ internodes of WT poplar plants under 6WL and 4WL+FR conditions. Bar, 50 μm. Right panel: Frequency distribution of the number of cells per cambium cell file presented in the left. One representative plant was shown. (**F**) Number of cells in each xylem cell file in the 7^th^ and 9^th^ internodes of WT poplar plants under 6WL and 4WL+FR conditions. Values are mean ± SD (n = 120). (**G**) Number of cells per cambium cell file in the 7^th^ and 9^th^ internodes of WT poplar plants under 6WL and 4WL+FR conditions. Values are mean ± SD (n = 120). Whiskers indicate the minimum and maximum values, and black lines within boxes indicate the median values. Different letters indicate significant differences (*P* < 0.05) by one-way ANOVA and LSD test for pairwise comparisons.

**Supplemental Figure S4.**
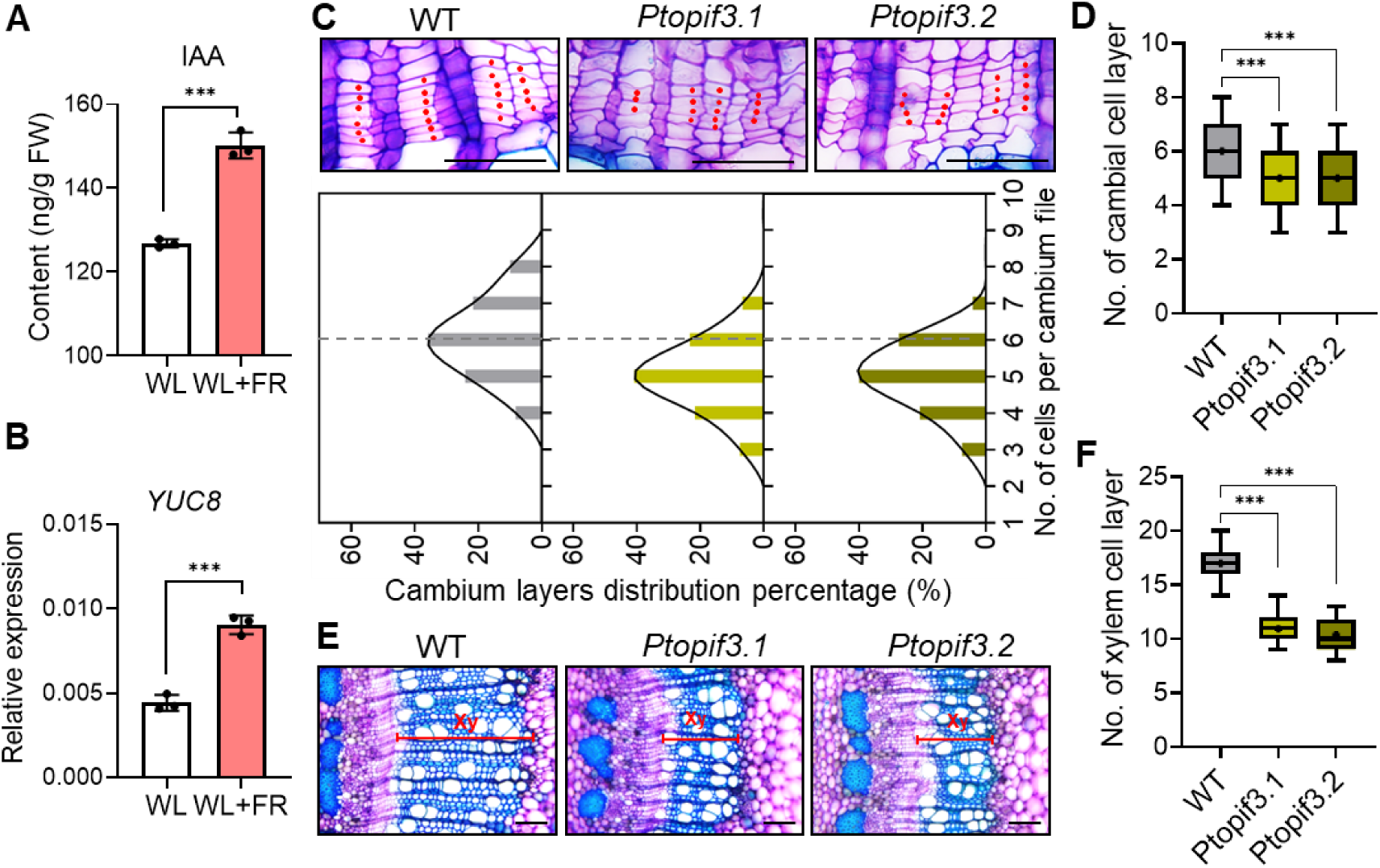
Vascular phenotypes of *Ptopif3.1 and Ptopif3.2* mutant plants. (**A**) Measurements of indole-3-acetic acid (IAA) content in the 6^th^ internodes of poplar WT plants under WL and shade (WL+FR) conditions Data are mean ± SD (n = 3 different plants). (**B**) qPCR analysis of auxin biosynthetic gene *YUC8* in the 6^th^ internodes of poplar WT plants under WL and shade (WL+FR) conditions. Data are mean ± SD (n = 3 different plants). (**C**) Close-ups of cambial cells in cross sections of the 7^th^ internodes from in WT, *Ptopif3.1* and *Ptopif3.2* transgenic plants (upper panel). Stem cross-sections were stained with Toluidine blue. Red dots indicate cambial cells in the same files. Bars, 20 μm. Frequency distribution of the number of cells per cambium cell file (lower panel). One representative plant was shown. (**D**) Number of cells per cambium cell file of WT, *Ptopif3.1* and *Ptopif3.2* transgenic plants. Values are mean ± SD (n = 120). (**E**) Xylem phenotypes in cross-sections of the 7^th^ internodes from in WT, *Ptopif3.1* and *Ptopif3.2* transgenic plants. Bars, 20 μm. (**F**) Number of xylem cell layers in the 7^th^ internodes of WT and *Ptopif3.1* and *Ptopif3.2* transgenic plants. Values are mean ± SD (n = 120). Whiskers indicate the minimum and maximum values, black lines within boxes denote the median values, with the means indicated by plus symbols. Asterisks denote significant differences by Student’s *t*-test (\*\*\**P* < 0.001).

**Supplemental Figure S5.**
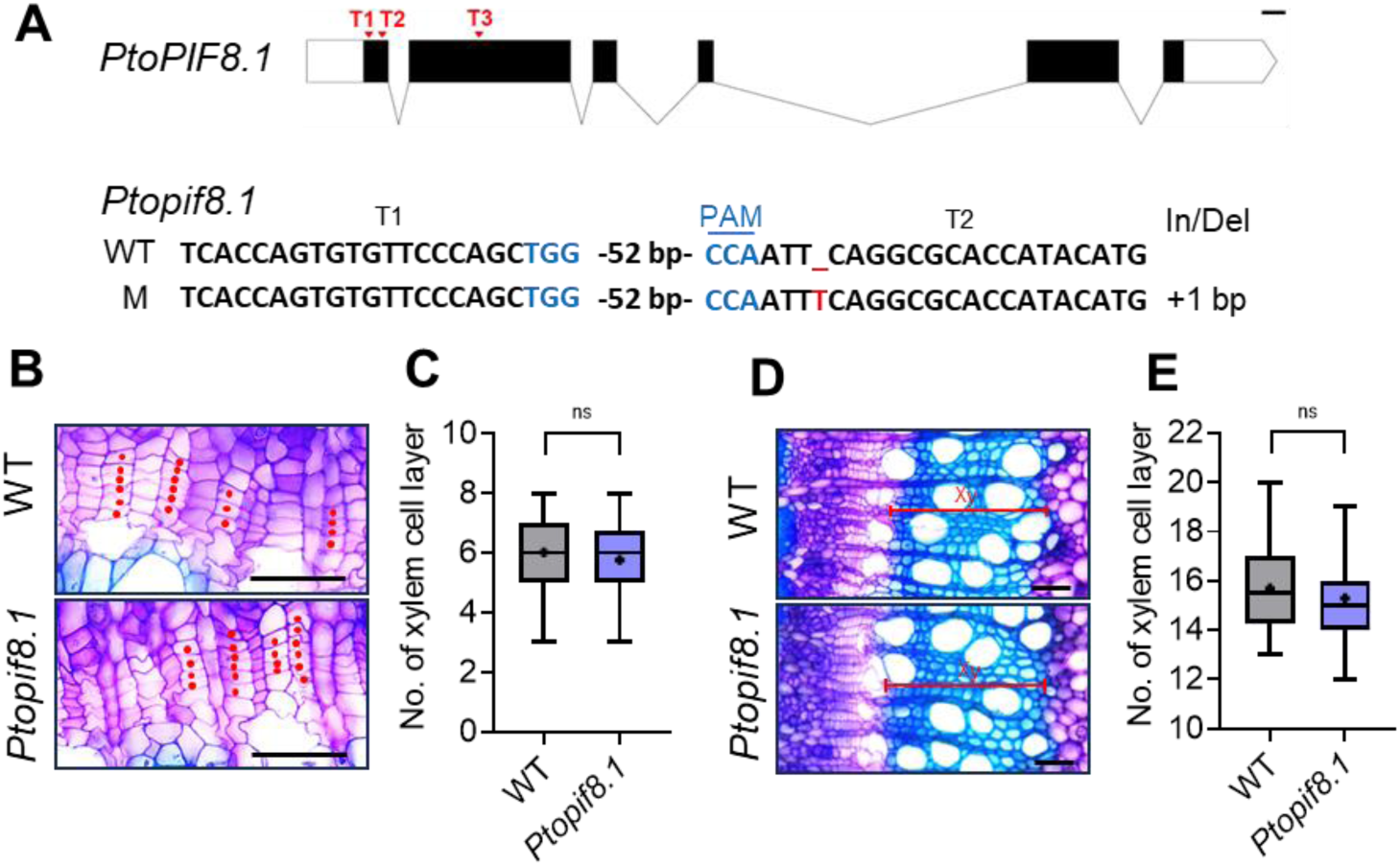
Vascular phenotypes of *Ptopif8.1* mutant plants. (**A**) Genotyping of *Ptopif8.1* single mutant. The gene structure of *PtoPIF8.1* with the locations of three single guide RNA targets (T1-T3) is present. Exons, introns, and the 5’-UTR/3’-UTR are denoted by black boxes, horizontal lines, and open boxes, respectively. Bar, 100 bp. PAM (NGG) sequences are in blue, and sequence insertion is indicated in red. (B) Cambium phenotypes in WT and *Ptopif8.1* transgenic plants. Red dots indicate cambial cells in the same files. Bars, 50 μm. (C) Statistical analysis of cambium cell layers in WT and *Ptopif8.1* transgenic plants. Values are mean ± SD (n = 120). (D) Xylem phenotypes in WT and *Ptopif8.1* transgenic plants. Bars, 100 μm. (E) Statistical analysis of xylem cell layers in WT and *Ptopif8.1* transgenic plants. Values are mean ± SD (n = 120). Whiskers indicate the minimum and maximum values, and black lines within boxes denote the median values, and the means are indicated by plus symbols. Not significant (ns) determined by Student’s *t*-test.

**Supplemental Figure S6.**
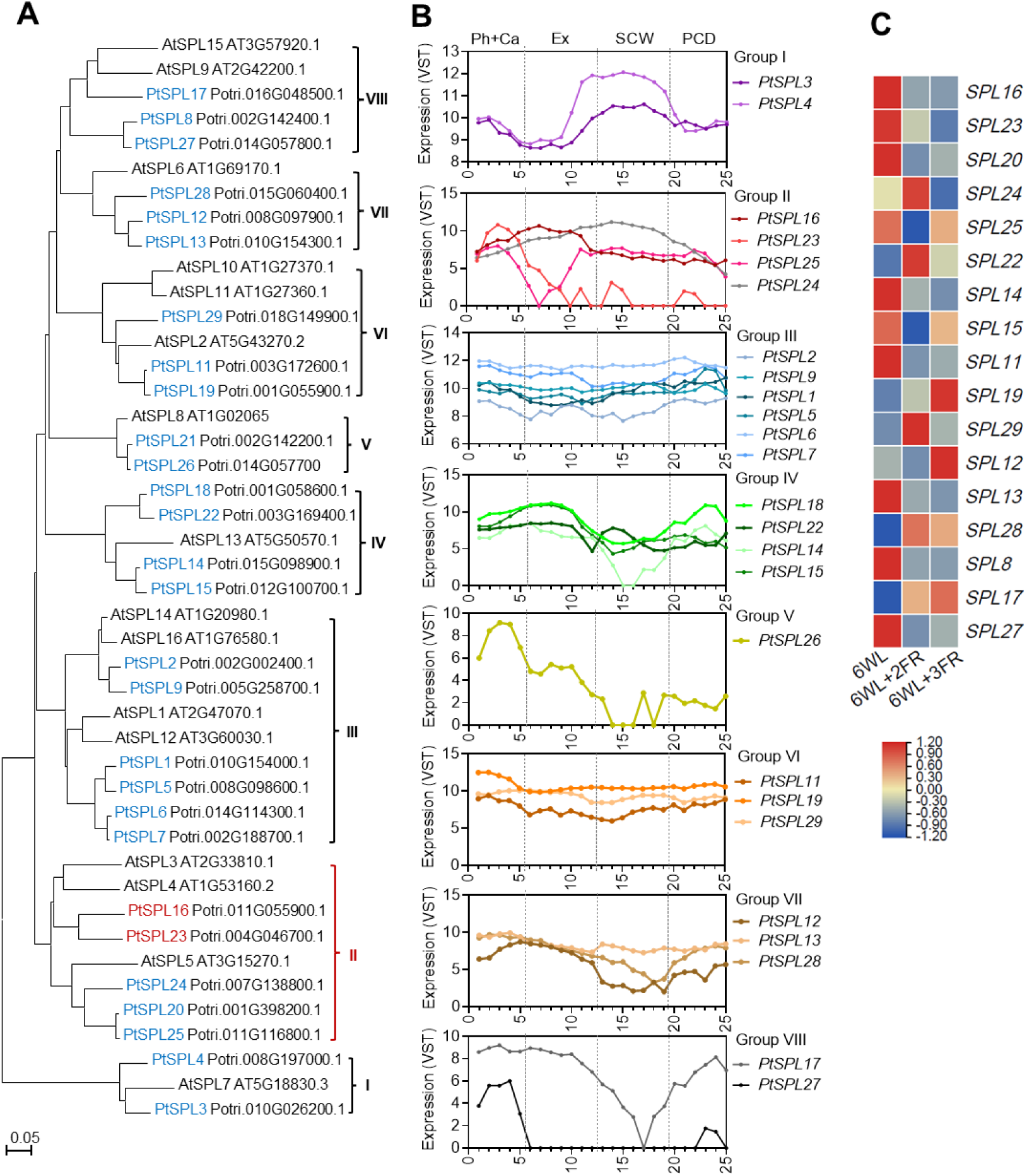
Expression profile of *Populus SPL* genes during wood development. (**A**) Phylogenetic analysis of *SPL* genes from *P. trichocarpa* (Pt) and *Arabidopsis thaliana* (At). Protein sequences were retrieved from the Phytozome database. Full-length of amino acid sequences were aligned with Clustal W using MEGA 6.0, and the phylogenetic tree was constructed using the neighbor-joining method. (**B**) Expression of *PtSPL* genes across the longitudinal wood sections of the aspen stem. Data were extracted from the AspWood database (Sundel et al., 2017). Data are shown across 25 sections from tree 1 (T1). Ph+Ca, phloem and cambium zone; Ex, expanding xylem; SCW; secondary cell wall forming xylem; PCD, programmed cell death zone. (**C**) Heatmap showing the expression changes of selected *Populus SPL* genes in the 7^th^ internodes of *P. tomentosa* WT plants grown under normal lights (6WL, R/FR = 10.69) and simulated shade conditions: 6WL+2FR (R/FR = 0.77) and 6WL+3FR (R/FR = 0.46). *SPL* genes specifically expressed in the phloem and cambium zones were selected for qPCR analysis in stem barks under normal light and simulated shade conditions. Data are mean ± SD (n = 3 different plants).

**Supplemental Figure S7.**
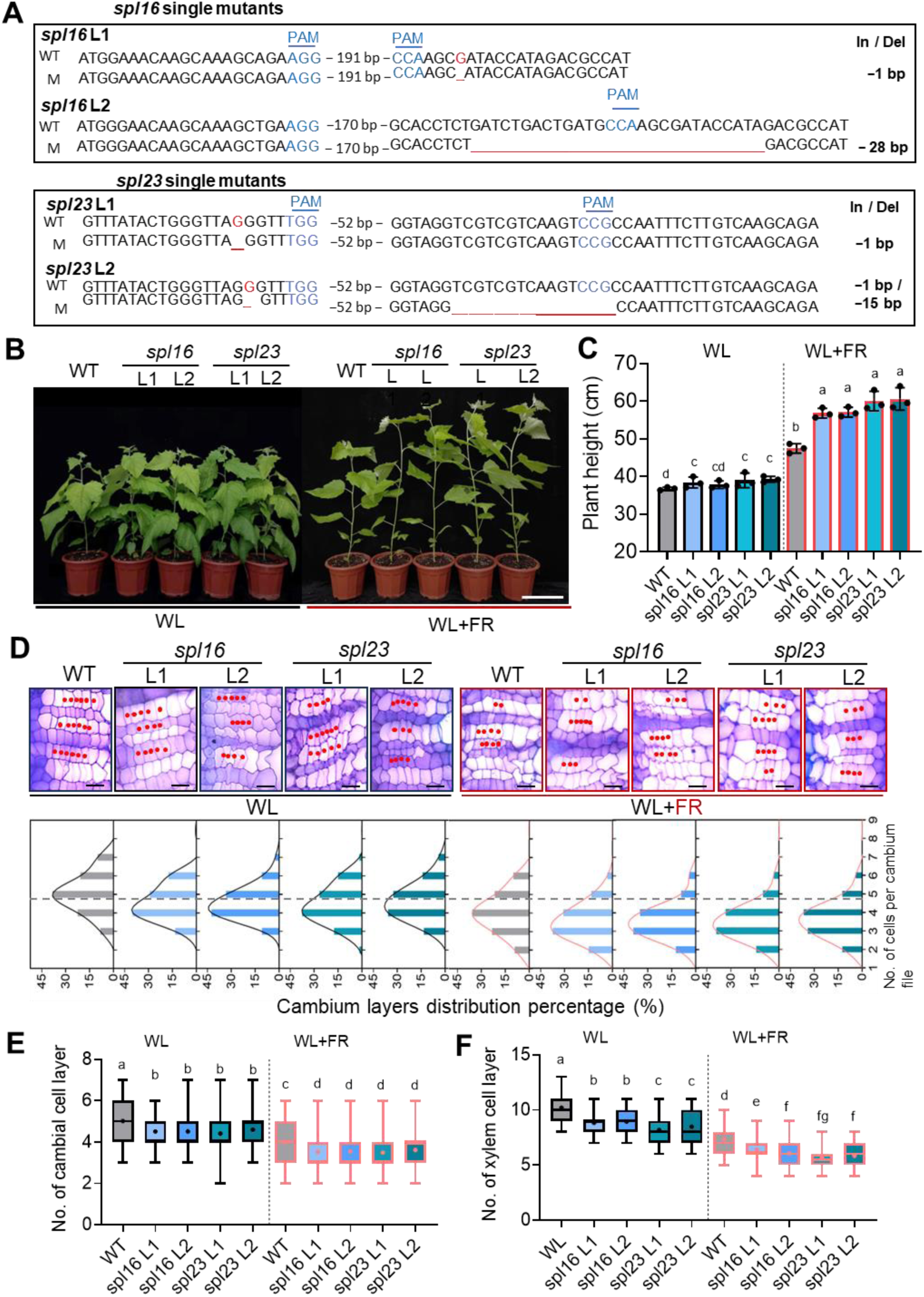
Cambial phenotypes of *P. tomentosa spl16* and *spl23* single mutants under normal light (WL) and simulated shade (WL+FR) conditions. (A) Genotyping of two independent lines of *P. tomentosa spl16* and *spl23* single mutants. The PAM (NGG) sequences were highlighted in blue. The *spl23* single mutants were generated in this study, while the *spl16* single mutants were produced in our previous study (Wei et al., 2023). The sequences in the CRISPR mutants were PCR-amplified, cloned, and sequenced for alignment with those in WT plants. (**B**) Growth phenotypes of *spl16*, *spl23* with WT poplars under WL and WL+FR conditions. Bar, 18 cm. (**C**) Plant height measurements of WT, *spl16* and *spl23* single mutant plants under WL and WL+FR conditions. Data are mean ± SD (n = 3 different plants). (**D**) Upper panel: Close-ups of cambial cells in cross-sections of the 7^th^ internodes from 6-week-old poplar WT, *spl16* and *spl23 s*ingle mutants. Stem cross-sections were stained with Toluidine blue. Cambial cells in each file were indicated by red dots. Bars, 20 μm. Lower panel: Frequency distribution of the number of cells per cambium cell file. One representative plant was shown. (**E**) Statistical analysis of the number of cells per cambial cell file of WT, *spl16* and *spl23* single mutants. Values are mean ± SD (n = 120). (**F**) Statistical analysis of xylem cell layers in the 7^th^ internode of WT, *spl16* and *spl23* single mutants. Values are mean ± SD (n = 130). Whiskers indicate the minimum and maximum values, black lines within boxes indicate the median values, and means are indicated by plus symbols. Different letters indicate significant differences (*P* < 0.05) by one-way ANOVA and LSD test for pairwise comparisons.

**Supplemental Figure S8.**
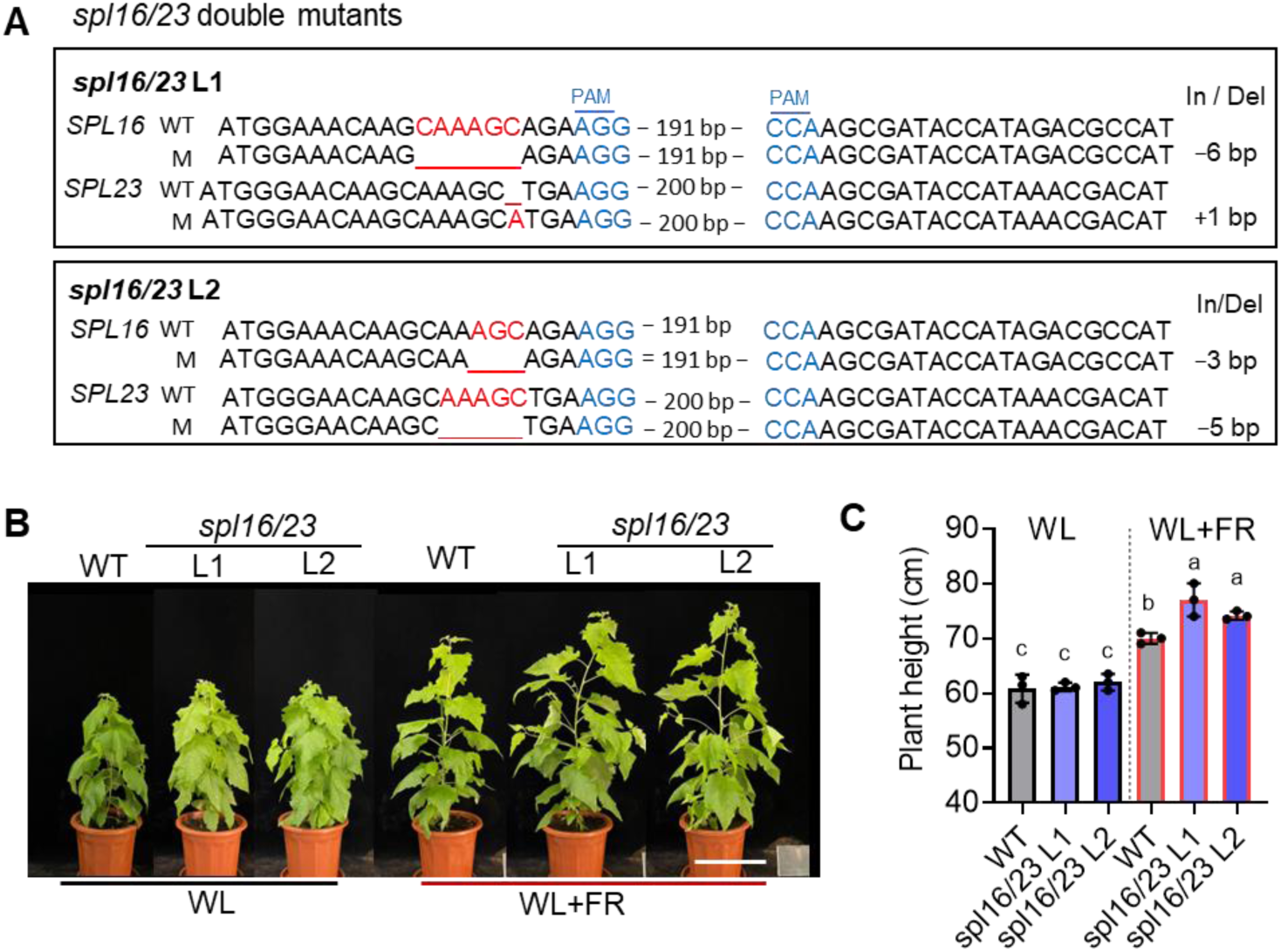
Phenotypes of the *spl16*/*23* double mutants under WL and WL+FR conditions. (**A**) The *P. tomentosa spl16*/*23* double mutants were generated in our previous study (Wei et al., 2023). The sequences in these mutants were PCR-amplified, cloned and sequenced for alignment with WT sequences. (**B**) Phenotypes of WT and *spl16/23* double mutants under WL and WL+FR conditions. Bar, 22 cm. (**C**) Measurements of plant height for WT and *spl16/23* double mutants under WL and WL+FR conditions. Data are mean ± SD (n = 3 different plants). Different letters indicate significant differences (*P* < 0.05) by one-way ANOVA and LSD test for pairwise comparisons.

**Supplemental Figure S9.**
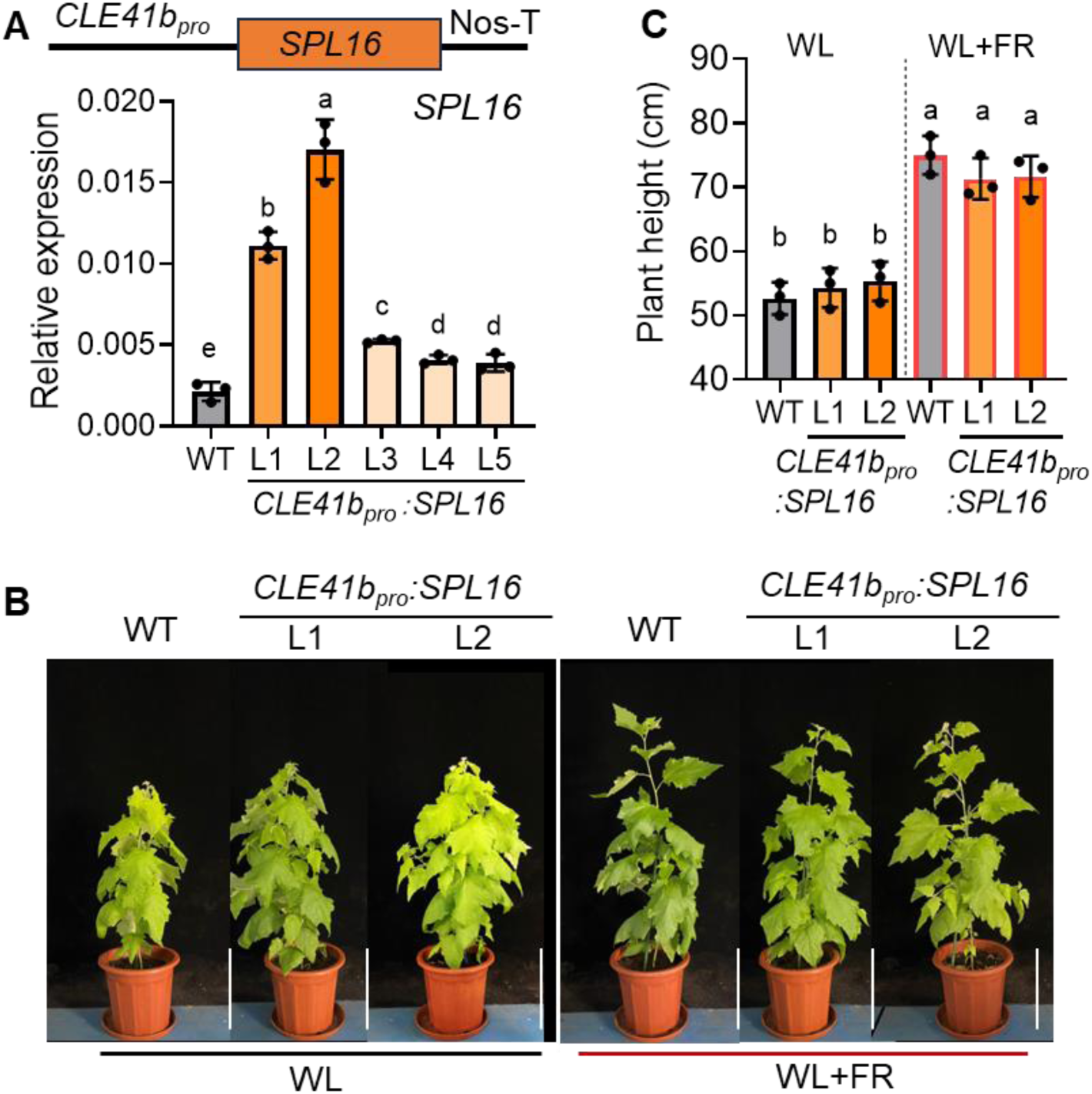
Generation of transgenic poplar with phloem-specific overexpression of *SPL16*. (**A**) Expression levels of *SPL16* in WT and *CLE41b_pro_:SPL16* lines, in which *SPL16* is driven by the phloem-specific promoter *CLE41b*. The miR156 target site is localized at the untranslated region (3’UTR) of *SPL16*, thus the *CLE41b_pro_:SPL16* construct is resistant to miR156. RNA was extracted from stem bark of the 7^th^ internode of WT and transgenic lines for qPCR analysis. Data are mean ± SD (n = 3 technical replicates). (**B**) Phenotypes of WT and *CLE41b_pro_:SPL16* transgenic plants under WL and WL+FR conditions. Bars, 22 cm. (**C**) Measurements of plant height for WT and *CLE41b_pro_:SPL16* transgenic plants under WL and WL+FR conditions. Data are mean ± SD (n = 3 different plants). Different letters indicate significant differences (*P* < 0.05) by one-way ANOVA and LSD test for pairwise comparisons.

**Supplemental Figure S10.**
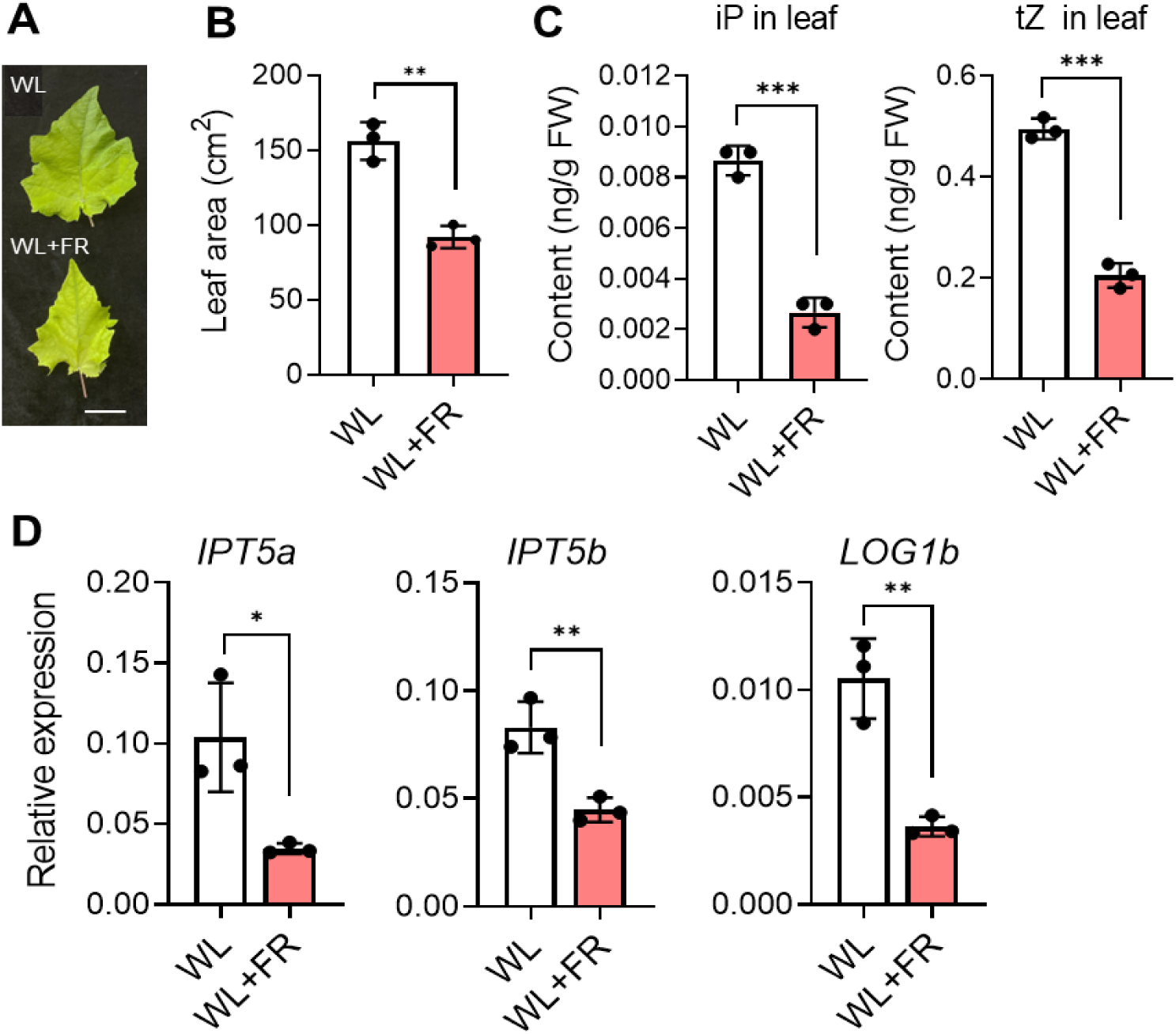
**Shade decreased leaf size and inhibited cytokinin levels in leaves.** (**A**) Comparison of leaf size from white light (WL)-grown and shade (WL+FR)-treated poplar plants. Representative mature leaf at the 7^th^ internode was shown. Bar, 2 cm. (**B**) Measurements of leaf area shown in (**A**). Data are mean ± SD from three different plants. (**C**) Quantification of endogenous CK contents in the leaves at the 8^th^ internode of poplar plants under WL and WL+FR conditions. Data are mean ± SD from three different plants. (**D**) qPCR analysis expression levels of CK biosynthesis genes (*IPT5a*, *IPT5b* and *LOG1b*) in mature leaves of poplar plants under WL and WL+FR conditions. Data are mean ± SD from three different plants. Asterisks denote significant differences by Student’s *t*-test (\**P* < 0.05; \*\**P* < 0.01; \*\*\**P* < 0.001).

**Supplemental Figure S11.**
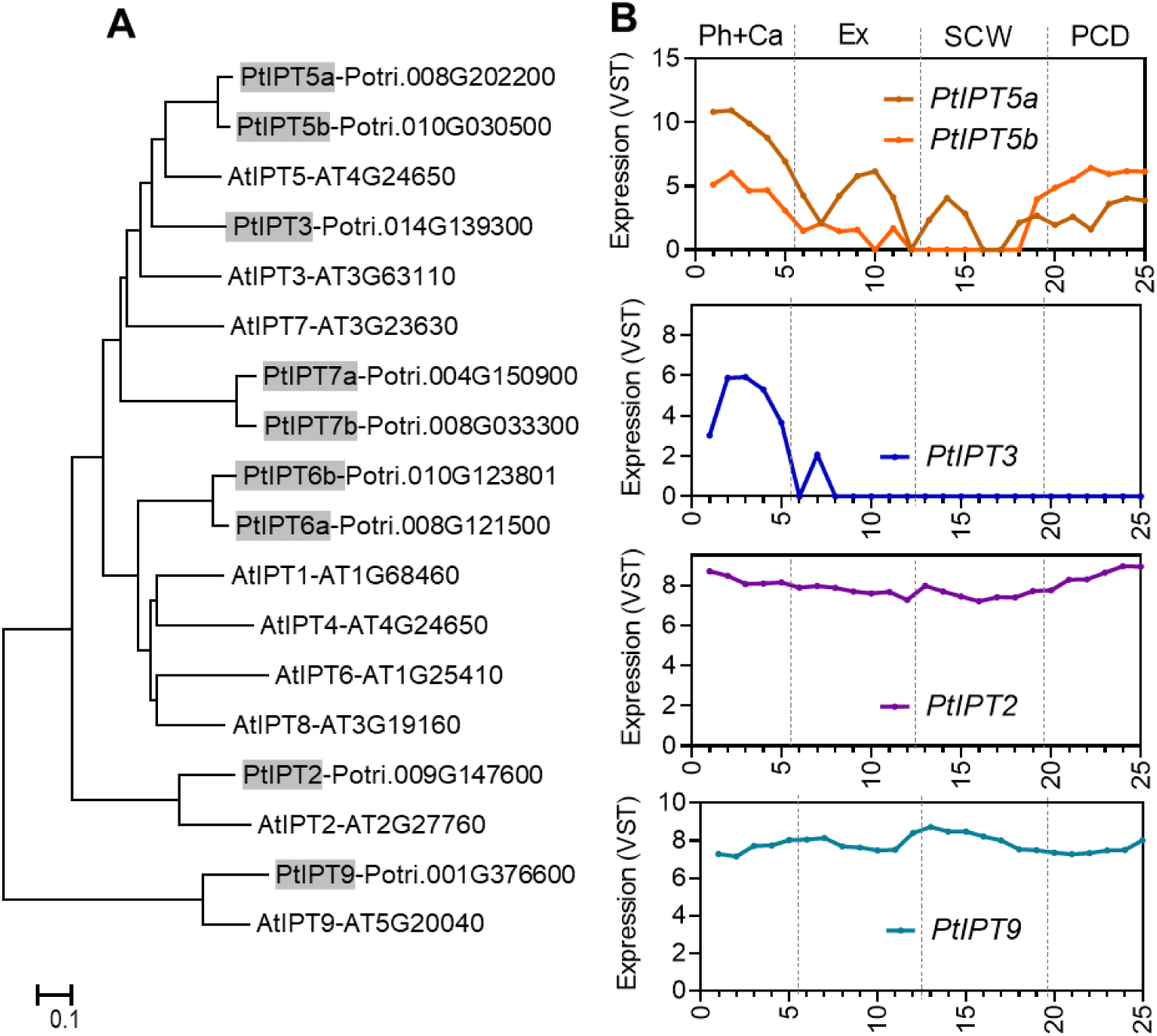
Expression profile of *Populus IPT* genes during wood development. (A) Phylogenetic analysis of *isopentenyltransferase* (*IPT*) genes from *P. trichocarpa* (Pt) and *A. thaliana* (At). Protein sequences were retrieved from the Phytozome database using given gene ID. Full-length of amino acid sequences were aligned with Clustal W using MEGA 6.0, and the phylogenetic tree was constructed using the neighbor-joining method. Bar = 0.1 substitutions per site. (B) Expression of *PtIPT* genes across the longitudinal wood sections of the aspen stem. Data were extracted from the AspWood database (Sundel et al., 2017). Data are shown across 25 sections from tree 1 (T1). Ph+Ca, phloem and cambium zone; Ex, expanding xylem; SCW; secondary cell wall forming xylem; PCD, programmed cell death zone.

**Supplemental Figure S12.**
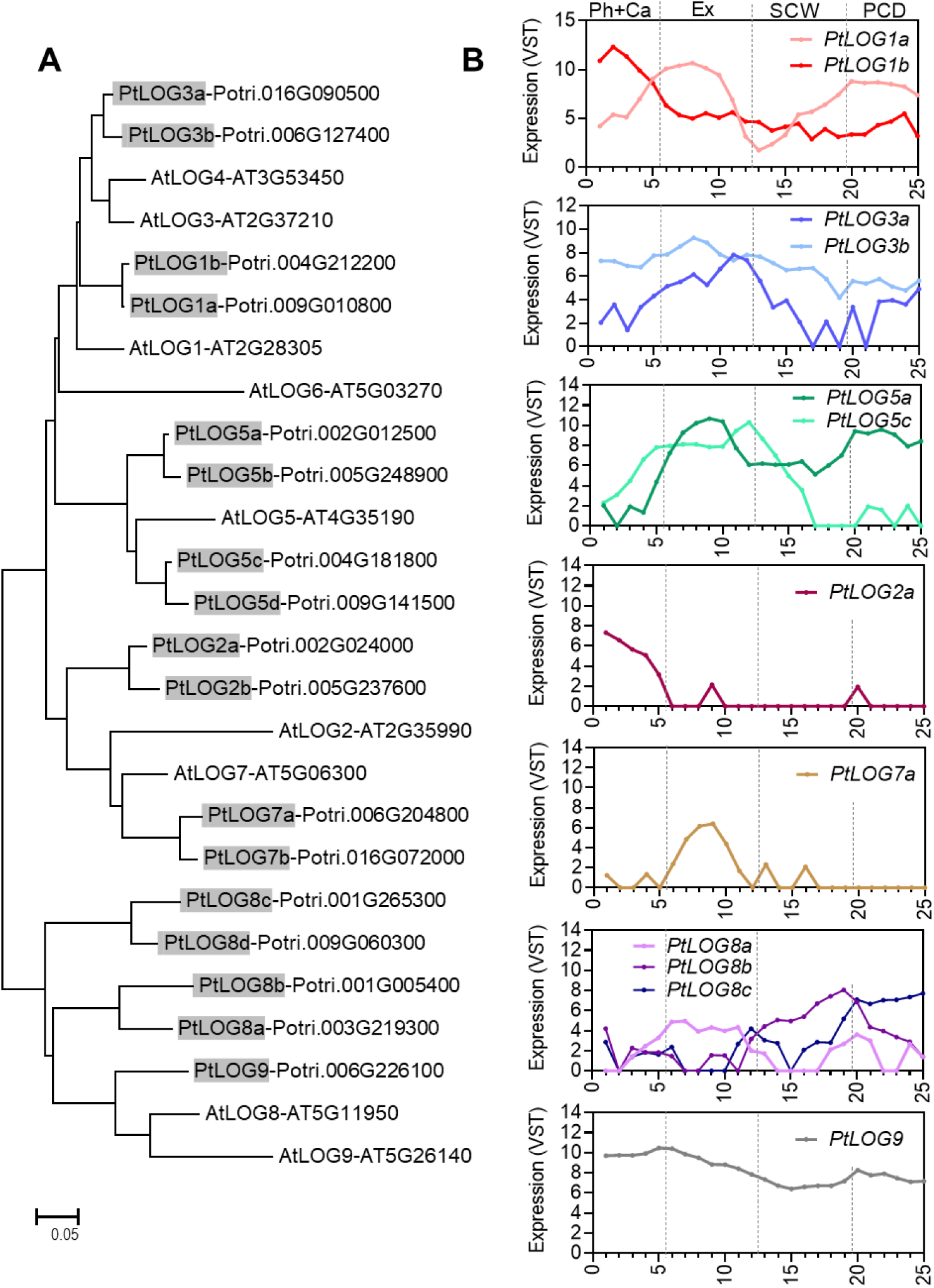
Expression profile of *Populus LOG* genes during wood development. (A) Phylogenetic analysis of *LONELY GUY* (*LOG*) genes from *P. trichocarpa* (Pt) and *A. thaliana* (At). *LOG* encodes a key enzyme that converts inactive cytokinin to biologically active form. Protein sequences were retrieved from the Phytozome database using given gene ID. Full-length of amino acid sequences were aligned with Clustal W using MEGA 6.0, and the phylogenetic tree was constructed using the neighbor-joining method. Bar = 0.1 substitutions per site. (**B**) Expression of *PtLOG* genes across the longitudinal wood sections of the aspen stem. Data were extracted from the AspWood database (Sundel et al., 2017). Data are shown across 25 sections from tree 1 (T1). Ph+Ca, phloem and cambium zone; Ex, expanding xylem; SCW; secondary cell wall forming xylem; PCD, programmed cell death zone.

**Supplemental Figure S13.**
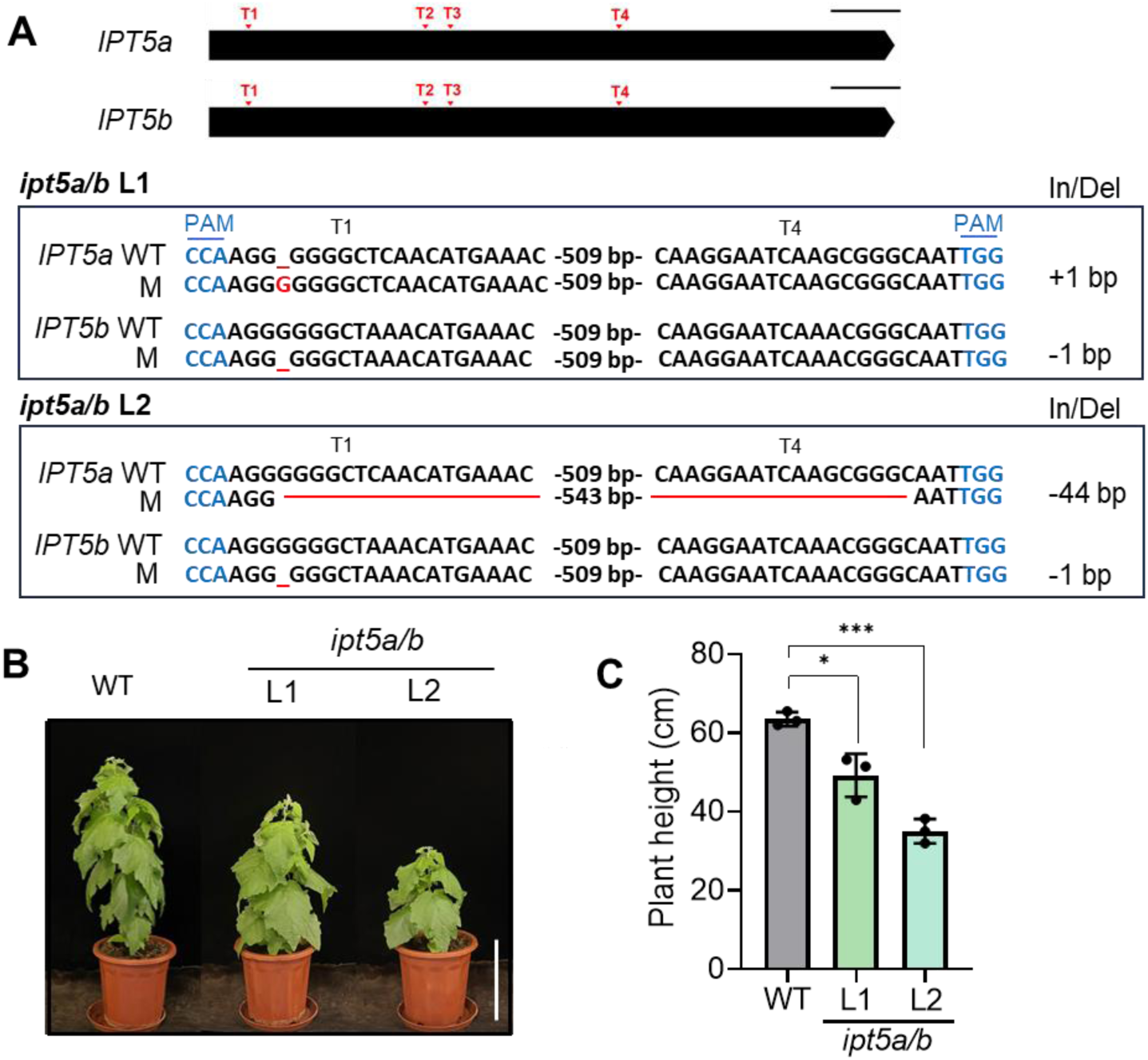
Generation of transgenic poplar with CRISPR/Cas9 knockout of *IPT5a/IPT5b*. (**A**) Genotyping of two independent lines of *P. tomentosa ipt5a/b* double mutants. Gene structure of *IPT5a* and *IPT5b* genes with the locations of four single guide RNA targets (T1-T4). Both *IPT5a* and *IPT5b* do not contain any intron. Bar, 100 bp. Genome editing occurred at the locations of target 1 and 4 sequences. The PAM (NGG) sequences were highlighted in blue. The sequence insertion or deletion was shown in red. (**B**) Phenotypes of WT and *ipt5a/b* transgenic plants. Bar, 11 cm. (**C**) Measurements of plant height for WT and *ipt5a/b* transgenic plants. Data are mean ± SD from three different plants. Asterisks indicate significant differences by Student’s *t*-test (*, *P* < 0.05; ***, *P* < 0.001).

**Supplemental Figure S14.**
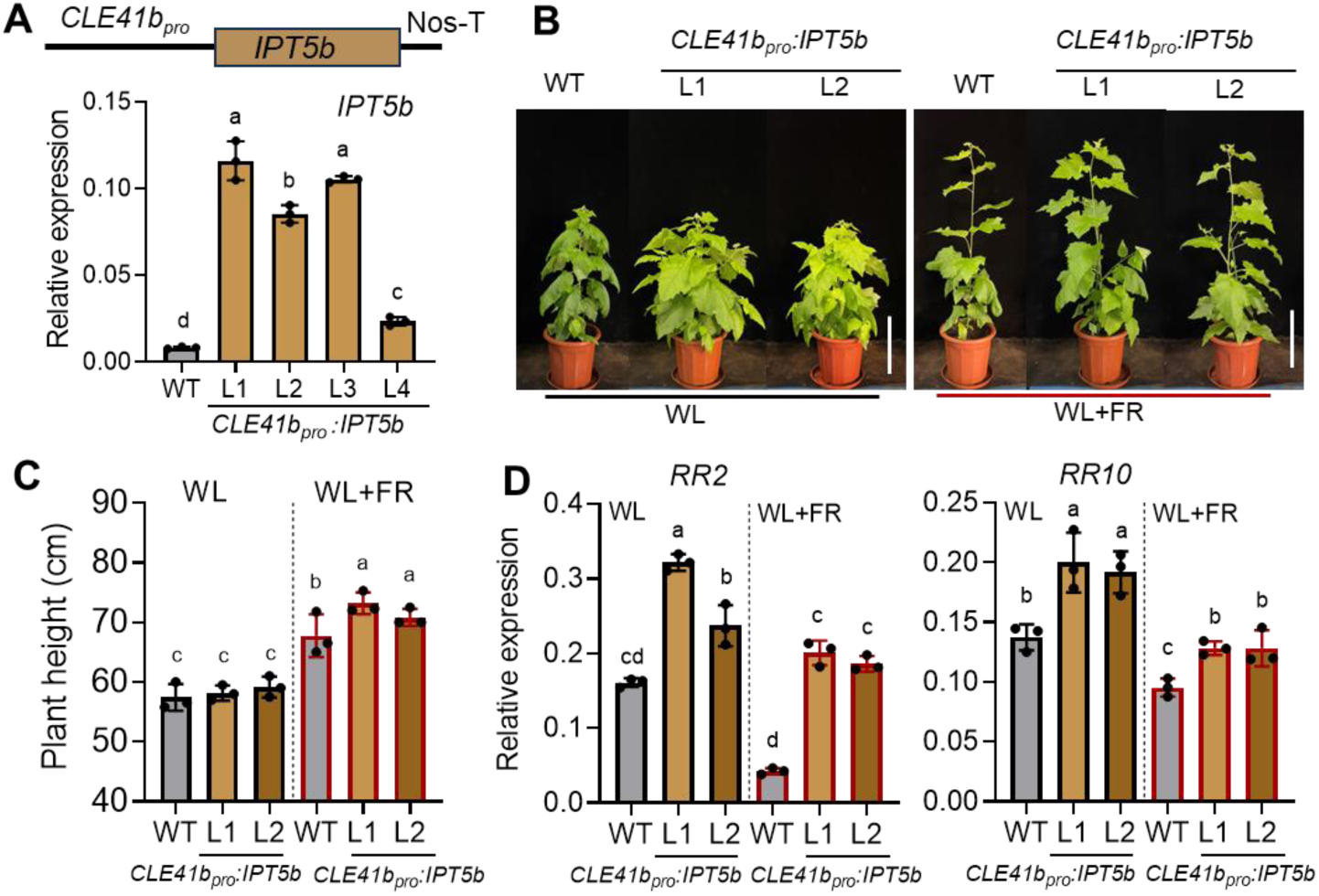
Generation of transgenic poplar plants with phloem-specific overexpression of *IPT5b*. (**A**) Expression levels of *IPT5b* in WT and *CLE41b_pro_:IPT5b* lines, in which *IPT5b* was driven by the phloem-specific promoter *CLE41b*. RNA was extracted from stem bark of the 7^th^ internode of WT and transgenic lines for qPCR analysis. Data are mean ± SD (n = 3 technical replicates). (**B**) Phenotypes of WT and *CLE41b_pro_:IPT5b* transgenic plants under normal WL and WL+FR conditions. Bar, 22 cm. (**C**) Measurements of plant height for WT and *CLE41b_pro_:IPT5b* transgenic lines under WL and WL+FR conditions. Data are mean ± SD (n = 3 different plants). (D) qPCR expression analysis of *RR2* and *RR10* in the stem bark of WT and *CLE41b_pro_:IPT5b* transgenic lines under WL and WL+FR conditions. Data are mean ± SD from three different plants. Different letters indicate significant differences (*P* < 0.05) by one-way ANOVA and LSD test for pairwise comparisons.

**Supplemental Figure S15.**
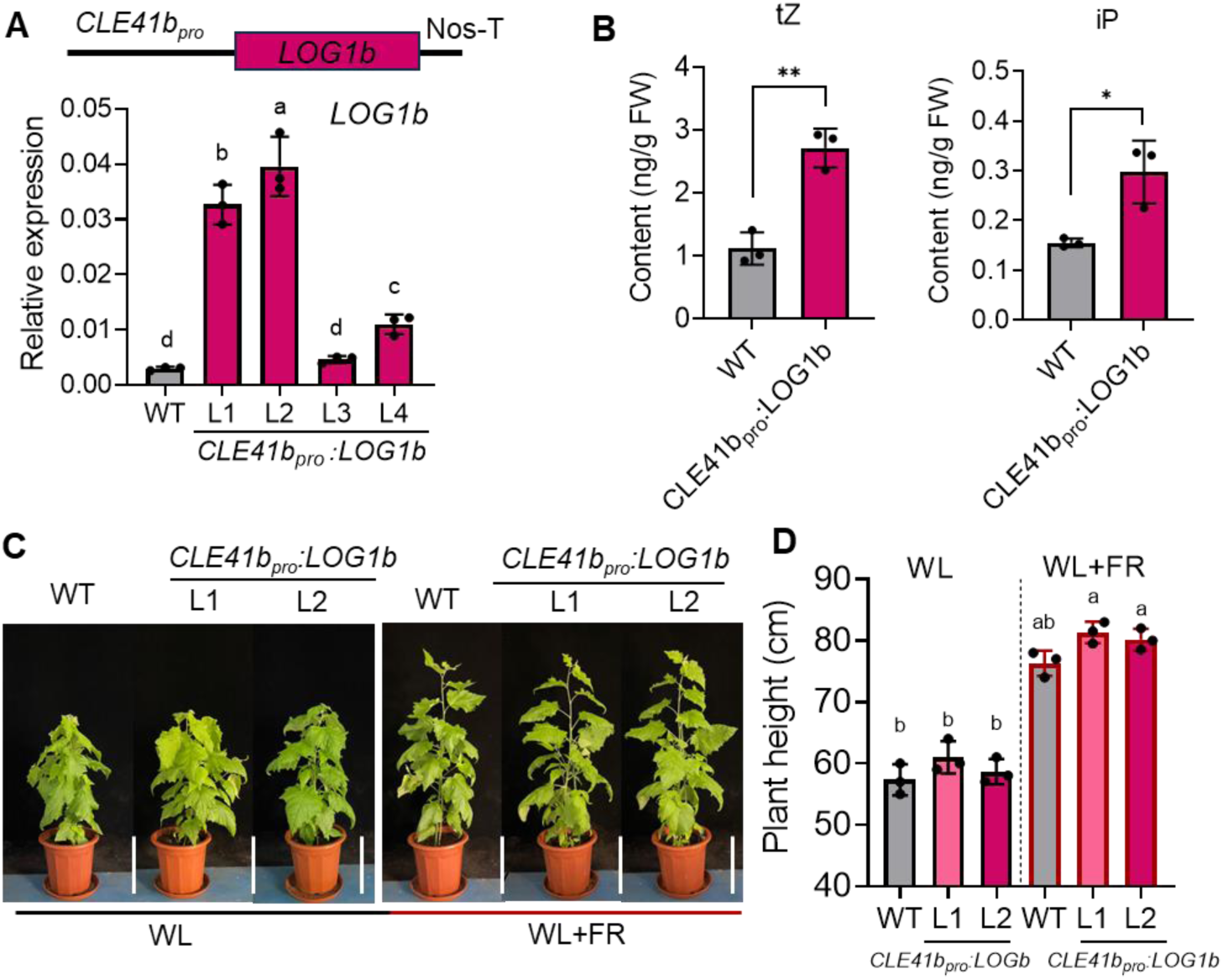
Generation of transgenic poplar plants with phloem-specific overexpression of *LOG1b*. (**A**) Expression levels of *LOG1b* in WT and *CLE41b_pro_:LOG1b* lines overexpressing *LOG1b* driven by the phloem-specific promoter *CLE41b*. RNA was extracted from stem bark of the 7^th^ internode of WT and transgenic lines for qPCR analysis. Data are mean ± SD (n = 3 technical replicates). (**B**) Quantification of tZ-type and iP-type CK content in the 6^th^ and 7^th^ internodes of WT and *CLE41b_pro_:LOG1b* (L2) transgenic plants grown under WL conditions. Data are mean ± SD from three different plants. Asterisks denote significant differences by Student’s *t*-test (\**P* < 0.05; \*\**P* < 0.01). (**C**) Phenotypes of poplar WT and *CLE41b_pro_:LOG1b* transgenic plants under WL and WL+FR conditions. Bars, 22 cm. (**D**) Measurements of plant height for WT and *CLE41b_pro_:LOG1b* transgenic lines under WL and WL+FR conditions. Data are mean ± SD from three different plants. Different letters in (A) and (D) indicate significant differences (*P* < 0.05) by one-way ANOVA and LSD test for pairwise comparisons.

**Supplemental Figure S16.**
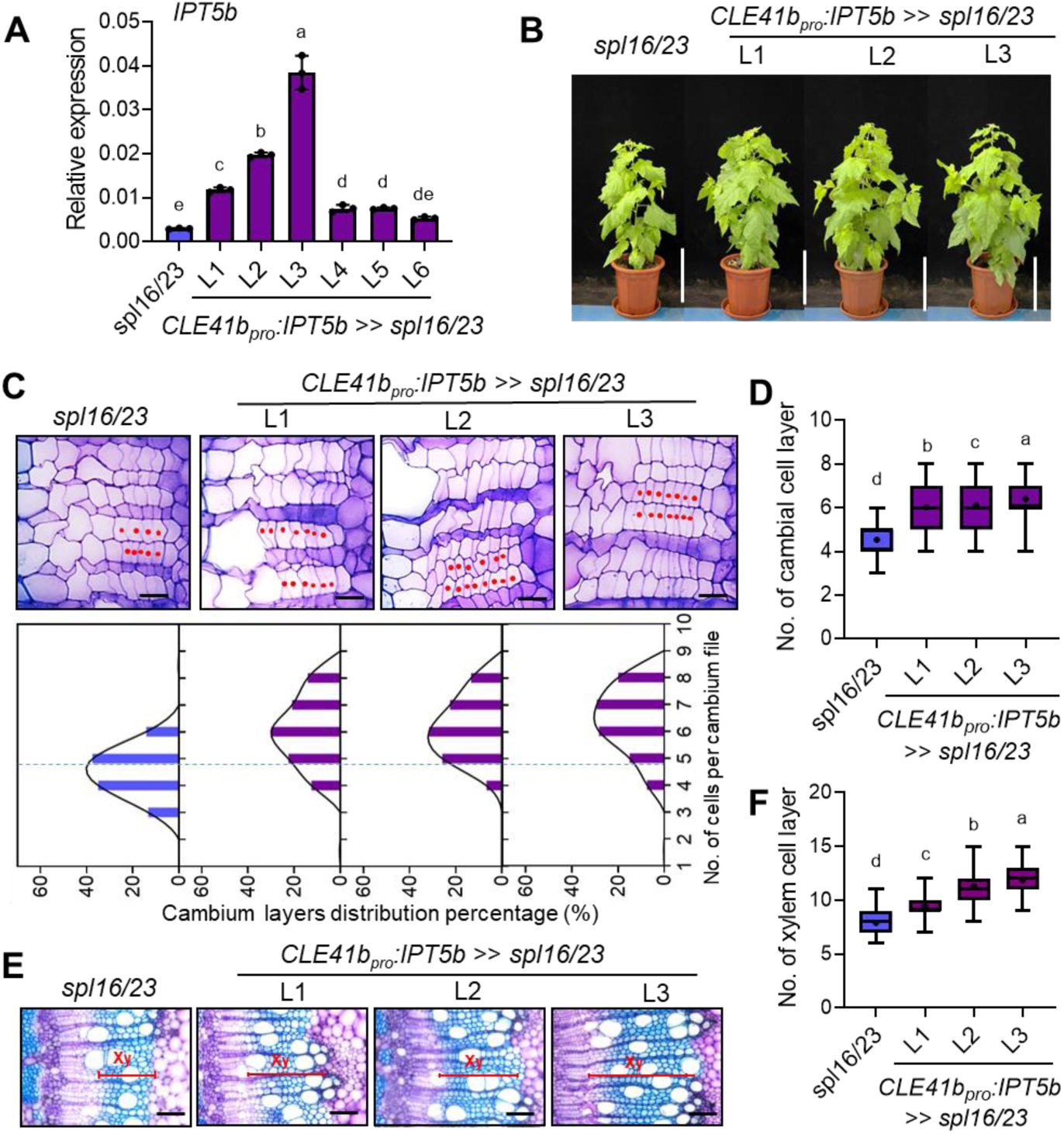
Overexpression of *IPT5b* rescues the vascular defect of *spl16/23* mutants. (**A**) Expression levels of *IPT5b* in *spl16/23* mutants and *CLE41b_pro_:IPT5b>>spl16/23* transgenic lines, in which *IPT5b* is overexpressed in the *spl16/23* mutant background. RNA was extracted from stem bark of the 7th internode of WT and transgenic lines for qPCR analysis. Values are mean ± SD (n = 3 technical replicates). (**B**) Phenotypes of *spl16/23* mutants and *CLE41b_pro_:IPT5b>>spl16/23* transgenic lines (L1-L3) under normal light for 8 weeks. Bar = 22 cm. (**C**) Close-ups of cambial cells in cross sections of the 7^th^ internodes from *spl16/23* mutant and *CLE41b_pro_:IPT5b>>spl16/23* transgenic lines (upper panel). Stem cross-sections were stained with Toluidine blue. Bars, 20 μm. Frequency distribution of the number of cells per cambium cell file (lower panel). One representative plant was shown. (**D**) Statistical analysis of the number of cells per cambium cell file of *spl16/23* mutant and *CLE41b_pro_:IPT5b>>spl16/23* transgenic lines. Values are mean ± SD (n = 120). (**E**) Xylem phenotypes in cross-sections of the 7^th^ internodes from *spl16/23* mutant and *CLE41b_pro_:IPT5b>>spl16/23* transgenic lines. Bars, 100 μm. (**F**) Statistical analysis of the number of xylem cell layers in the 7^th^ internode of *spl16/23* mutant and *CLE41b_pro_:IPT5b>>spl16/23* transgenic lines. Values are mean ± SD (n = 118). Whiskers indicate the minimum and maximum values, black lines within boxes denote the median values, and means are indicated by plus symbols. Different letters indicate significant differences (*P* < 0.05) by one-way ANOVA and LSD test for pairwise comparisons.

**Supplemental Figure S17.**
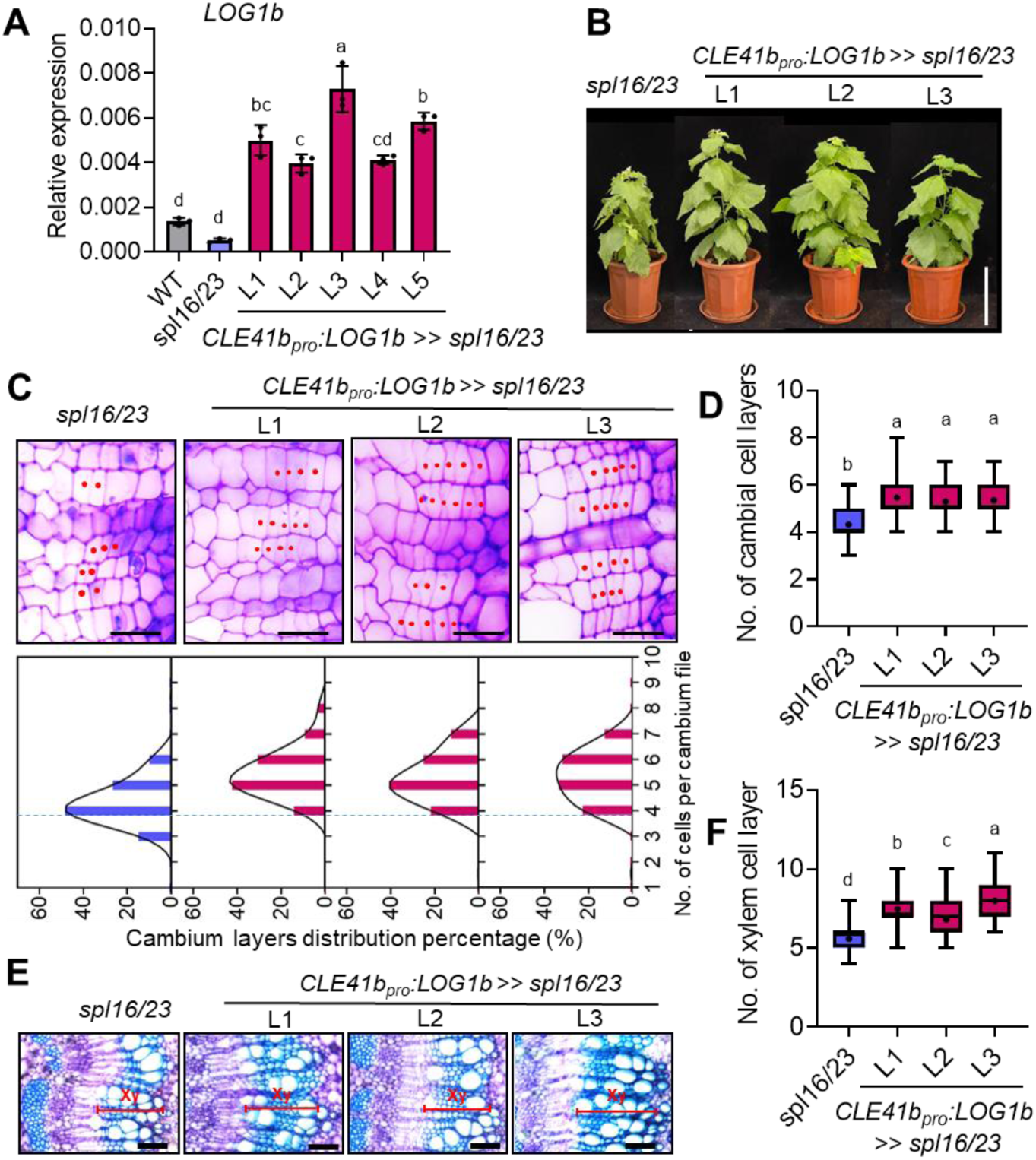
Overexpression of *LOG1b* rescues the vascular defect of *spl16/23* mutants. **(A)** Expression levels of *LOG1b* in WT, *spl16/23* mutant, and *CLE41b_pro_:LOG1b>>spl16/23* transgenic lines, in which *LOG1b* is overexpressed in the *spl16/23* mutant background. RNA was extracted from stem bark of the 7^th^ internode of different genotypes for qPCR analysis. Values are mean ± SD (n = 3 technical replicates). (**B**) Phenotypes of *spl16/23* mutants and *CLE41b_pro_:LOG1b>>spl16/23* transgenic lines grown under normal light for 6 weeks. Bar = 22 cm. (**C**) Close-ups of cambial cells in cross sections of the 7^th^ internodes from the *spl16/23* mutant and *CLE41b_pro_:LOG1b>>spl16/23* transgenic lines (upper panel). Stem cross-sections were stained with toluidine blue. Bars, 20 μm. Frequency distribution of the number of cells per cambium cell file (lower panel). One representative plant was shown. (**D**) Statistical analysis of the number of cells per cambium cell file of *spl16/23* mutant and *CLE41b_pro_:LOG1b>>spl16/23* transgenic lines. Values are mean ± SD (n = 120). (**E**) Xylem phenotypes in cross-sections of the 7^th^ internodes from *spl16/23* mutant and *CLE41b_pro_:LOG1b>>spl16/23* transgenic lines. Bars, 100 μm. (**F**) Statistical analysis of the number of xylem cell layers in the 7^th^ internode of *spl16/23* mutant and *CLE41b_pro_:LOG1b>>spl16/23* transgenic lines. Values are mean ± SD (n = 120). Whiskers indicate the minimum and maximum values, black lines within boxes indicate the median values, and means are indicated by plus symbols. Different letters indicate significant differences (*P* < 0.05) by one-way ANOVA and LSD test for pairwise comparisons.

